# Unrestrained fatty acid oxidation triggers heart failure in mice via cardiolipin loss and mitochondrial dysfunction

**DOI:** 10.64898/2026.01.12.698694

**Authors:** Chai-Wan Kim, Goncalo Vale, Xiaorong Fu, Jeffrey McDonald, Chongshan Dai, Chao Li, Zhao V. Wang, Gaurav Sharma, Chalermchai Khemtong, Craig Malloy, Stanislaw Deja, Shawn C. Burgess, Matthew Mitsche, Jay D. Horton

## Abstract

Cardiomyocytes primarily rely on fatty acid oxidation (FAO), which provides more than 70% of their energy. However, excessive FAO can disrupt cardiac metabolism by increasing oxygen demand and suppressing glucose utilization through the Randle cycle. Although inhibition of FAO has been investigated in heart failure, its overall therapeutic impact remains uncertain. To determine the consequences of enhanced FAO, we generated cardiomyocyte-specific ACC1 and ACC2 double-knockout (ACC dHKO) mice, which exhibit constitutively elevated FAO. ACC dHKO mice developed dilated cardiomyopathy and heart failure. Lipidomic analysis revealed marked depletion of cardiolipin caused by reduced linoleic acid, a direct consequence of excessive FAO. This cardiolipin deficiency impaired mitochondrial electron transport chain (ETC) activity, leading to mitochondrial dysfunction. Pharmacologic inhibition of FAO with etomoxir or oxfenicine restored cardiolipin levels, normalized ETC activity, and prevented cardiac dysfunction in ACC dHKO mice. These findings demonstrate that unrestrained FAO disrupts both lipid and energy homeostasis, culminating in heart failure in this model. Collectively, these results indicate that although FAO is essential for cardiac energy production, therapeutic strategies aimed at stimulating cardiac FAO may be detrimental rather than beneficial in heart failure.

## Introduction

Cardiomyopathy is a prevalent and debilitating form of heart failure that encompasses a diverse group of myocardial disorders characterized by structural and functional abnormalities, often leading to compromised cardiac function and, in many cases, heart failure (1–3). Currently, heart failure affects approximately 6.2 million people in the U.S. (4), with cardiomyopathy being a leading underlying cause (5). Moreover, the incidence of cardiomyopathy is rising, driven by various risk factors, including obesity, hypertension, and diabetes (6–8).

Recent studies have emphasized the pivotal role that alterations in cardiac energy metabolism play in the pathogenesis of cardiomyopathy (see (9, 10) for reviews). Remarkably, the human heart consumes ∼6 kg of ATP daily. More than 95% of this ATP is supplied through the oxidative phosphorylation of fatty acids (FAs) and glucose (2, 11). In a healthy heart, 60-90% of ATP is generated through FA oxidation while the contribution of glucose oxidation is 10-40% (10). However, in heart failure, the proportion of FA oxidation decreases relative to glucose utilization as the disease progresses (1, 2). Since FA oxidation requires 10-15% more oxygen than glucose oxidation, the decrease in FA oxidation may be a compensatory mechanism of the failing heart to conserve oxygen (12, 13). Although there is substantial debate as to whether changes in the ratio of FA and glucose oxidation are causally linked to the development of heart failure (10, 13, 14), several clinical studies have suggested that partial inhibition of FA oxidation has therapeutic potential for heart failure (15, 16).

In recent decades, insights into the intricate interplay between FA and glucose oxidation and its profound impact on the development of cardiomyopathy have been revealed (10). A key regulatory enzyme that determines substrate utilization within the heart is acetyl-CoA carboxylase (ACC). ACC is responsible for the conversion of acetyl-CoA to malonyl-CoA, a molecule with dual functions (17). Malonyl-CoA serves as a building block for FA synthesis (18) and as a regulator of FA oxidation by inhibiting carnitine palmitoyltransferase 1 (CPT1), the transporter of long chain FAs into mitochondria (19). By modulating FA oxidation, ACC influences the balance between FA and glucose oxidation. All cells express two forms of ACC. ACC1 is in the cytosol and produces malonyl-CoA for FA synthesis. ACC2 is associated with the mitochondrial membrane and produces malonyl-CoA primarily to regulate CPT1 (20). Cardiomyocytes have very low rates of *de novo* lipogenesis, and the expression of ACC1 is very low compared to lipogenic organs such as the liver (20, 21). In contrast, the expression of ACC2 in cardiomyocytes is very high compared to most other tissues (21).

Mouse models with altered ACC expression in various tissues have provided compelling evidence that dysregulation of this enzyme is associated with changes in FA and glucose oxidation (18, 22, 23). A complete understanding of the intricate role of ACCs in cardiac energy metabolism may open new avenues for developing tailored approaches to prevent or manage cardiomyopathy and improve patient outcomes.

In this study, deleting ACC1 and ACC2 in cardiomyocytes led to unrestrained FA oxidation and a significant reduction of cardiolipin, causing mitochondrial dysfunction and ultimately heart failure. These deleterious effects were prevented in mice lacking ACC1 and ACC2 by administering drugs that suppress FA oxidation. These findings highlight the potential risk of elevated FAO and urge caution in adapting strategies aimed at promoting FA oxidation in failing hearts.

## Results

Cardiomyocyte-specific ACC1 (ACC1 HKO), ACC2 (ACC2 HKO), and double-knockout (ACC dHKO) mice were generated by crossing *Acc1*– and *Acc2*-floxed alleles with *Myh6*-Cre transgenic mice (18). Deletions of *Acc1* and *Acc2* were verified by quantitative PCR (qPCR) (Figure 1A and B). *Acc1* mRNA levels were reduced by ∼60% in ACC1 HKO and ACC dHKO hearts, while *Acc2* mRNA levels were reduced by ∼95% in ACC2 HKO and ACC dHKO hearts. As indicated by qPCR Ct values, basal *Acc1* expression in heart was substantially lower than *Acc2*, consistent with prior reports (24). The residual *Acc1* and *Acc2* mRNA detected in the respective cardiomyocyte-specific knockouts was likely due to non-cardiomyocyte expression within whole-heart RNA preparations. To confirm functional loss of ACC activity, we measured malonyl-CoA, the enzymatic product of ACC (Figure 1C). Total malonyl-CoA concentrations were ∼89% lower in ACC dHKO hearts than in WT controls, whereas acetyl-CoA (the substrate of ACC) was unchanged across genotypes.

**Figure 1.**
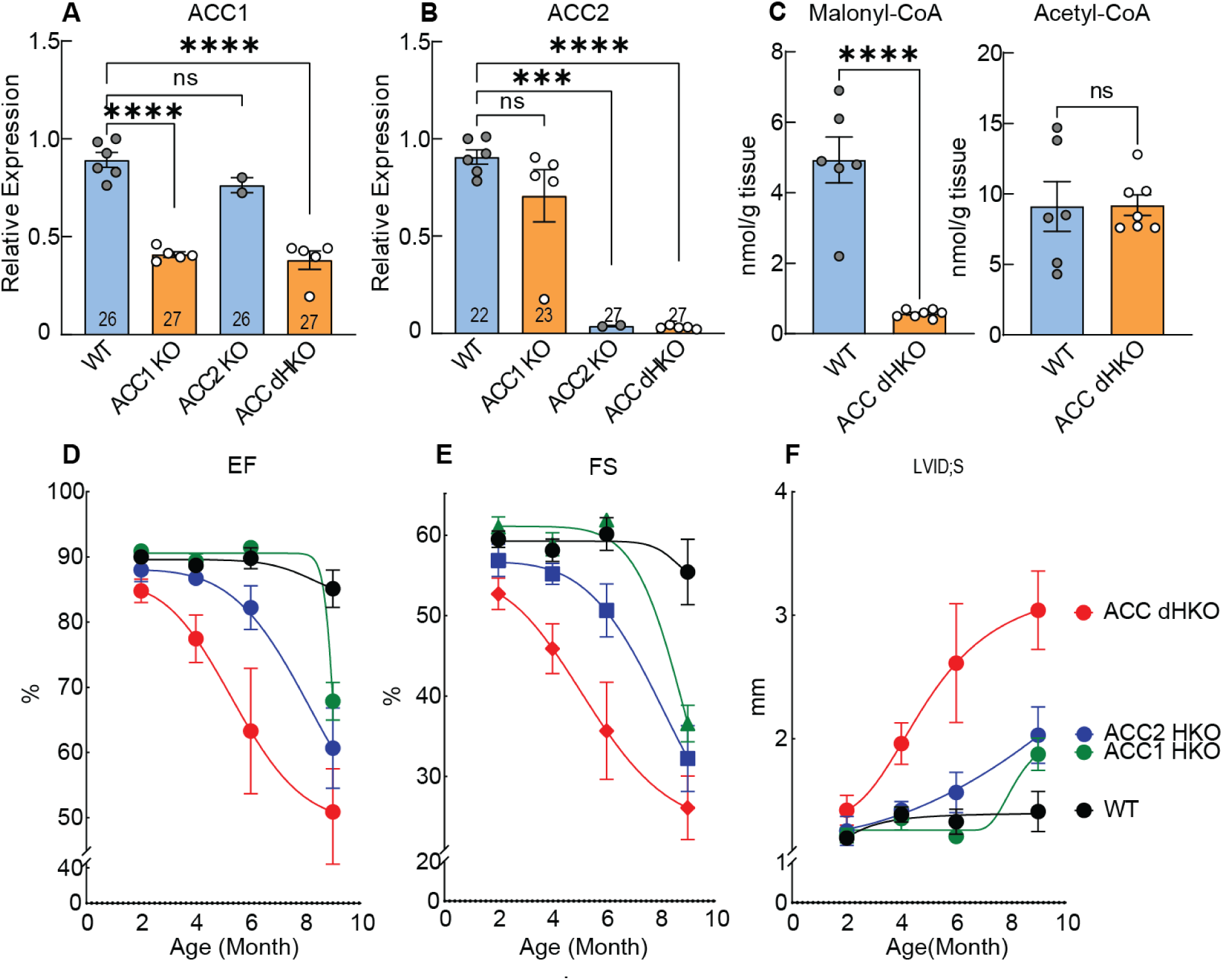
Genetic deletion of ACC1 and ACC2 in mouse cardiomyocytes. (**A** and **B**) Total RNA was isolated from hearts of 10-week-old WT, ACC1 HKO, ACC2 HKO, and ACC dHKO male mice, and quantitative RT-qPCR was performed to measure *Acc1* and *Acc2* mRNA expression. ***P < 0.001, ****P < 0.0001 by one-way ANOVA. (**C**) Hearts from 20-week-old WT and ACC dHKO female mice were harvested and freeze-clamped to measure malonyl-CoA and acetyl-CoA levels as described in Methods. ****P < 0.0001 by unpaired two-tailed Student’s t test. (**D–F**) ECHO assessment of cardiac function in male ACC1 HKO, ACC2 HKO, and ACC dHKO mice at 2, 4, 6, and 9 months of age.

Cardiac function was assessed by echocardiography (ECHO). ACC1 HKO mice exhibited normal cardiac function up to ∼6 months, but by 9 months exhibited a 20% reduction in ejection fraction (EF) and a 34% reduction in fractional shortening (FS) relative to WT (Figure 1D–E). ACC2 HKO mice developed mild dysfunction by 6 months (∼10% decrease in EF and FS) that progressed to 29% and 42% reductions in EF and FS, respectively, by 9 months (Figure 1D–E). In contrast, ACC dHKO mice exhibited systolic dysfunction by 2 months, with increased left ventricular internal diameter (LVID) during systole and reduced EF and FS (Figure 1F). Cardiac function then progressively declined, with EF and FS falling to 40% and 53% of WT, respectively, by 9 months. Together, these data demonstrate that combined loss of ACC1 and ACC2 in cardiomyocytes leads to early-onset and progressive cardiomyopathy in this model, whereas deletion of either isoform alone results in delayed or milder impairment.

Given the potential functional overlap between ACC1 and ACC2 in cardiomyocytes, we further characterized ACC double-knockout (ACC dHKO) mice to define mechanisms leading to heart failure in this model. Hematoxylin and eosin (H&E) staining revealed marked left ventricular enlargement in 11-month-old ACC dHKO hearts compared to WT controls (Figure 2A). Consistent with this, ECHO demonstrated increased LVID during both systole and diastole, accompanied by reduced left ventricular posterior wall (LVPW) thickness (Figure 2B–C). Systolic performance was markedly impaired in ACC dHKO mice, with an ∼51% reduction in EF and ∼62% reduction in FS at 10 months relative to WT (Figure 2C). These functional deficits were associated with increased expression of fetal cardiac genes *Nppa*, *Nppb*, and *Myh7*, molecular markers of pathological hypertrophy and heart failure (Figure 2D) (25, 26). Masson’s trichrome staining revealed extensive interstitial fibrosis, with a fibrotic area >5-fold greater in ACC dHKO hearts than in WT hearts (Figure 2E). Pulmonary congestion was also assessed in ACC dHKO mice. Lung fluid weight correlated with cardiac dysfunction (EF and FS) in ACC dHKO mice (Supplemental Figure 1A) (27), consistent with pulmonary edema accompanying progressive cardiac decompensation. However, lung weight–to–body weight (LW/BW) ratios did not differ between WT and ACC dHKO mice (Supplemental Figure 1B).

**Figure 2.**
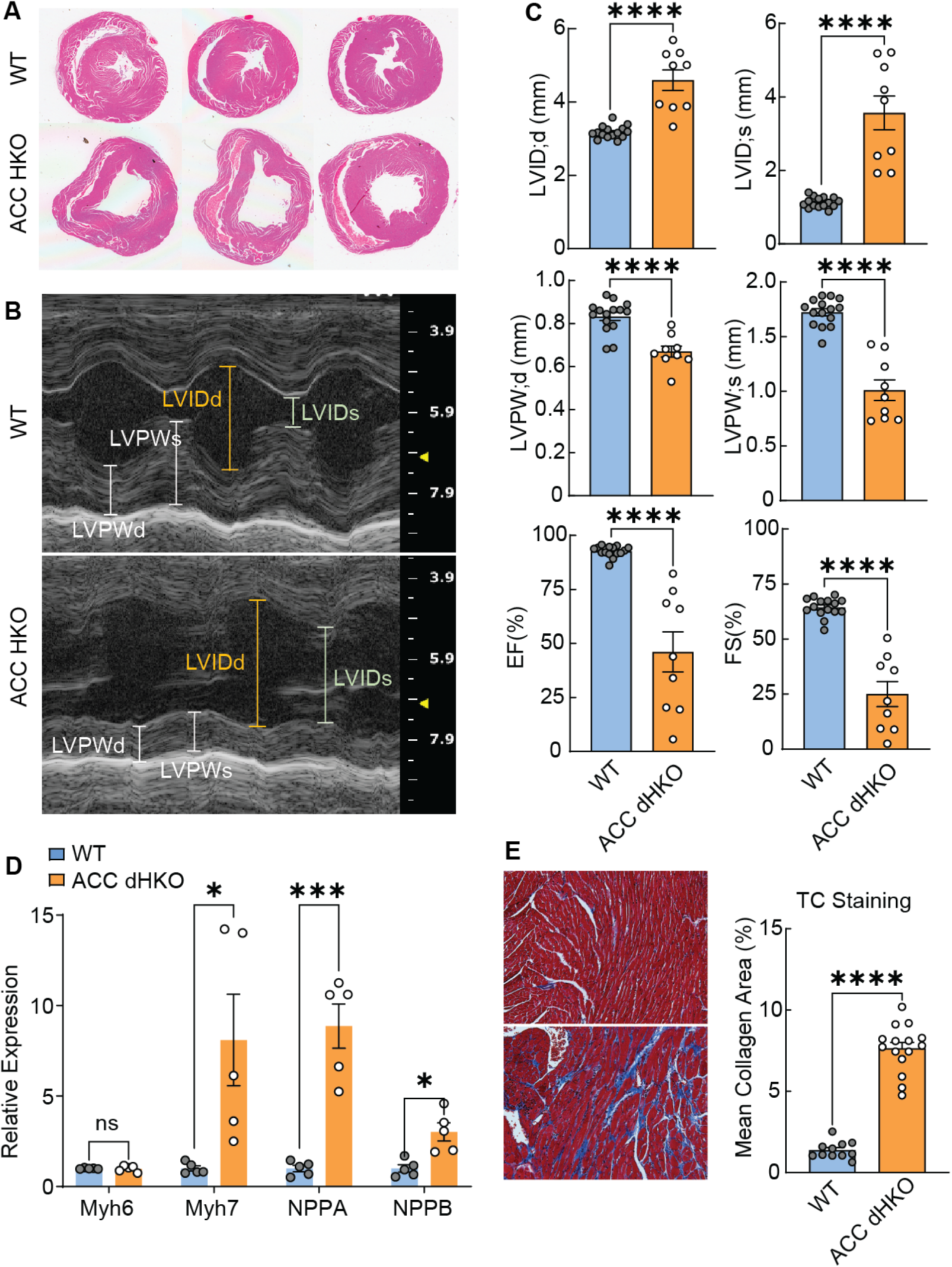
Decreased heart function in ACC dHKO mice. (**A**) Transverse/two-chamber views of hearts from 11-month-old WT and ACC dHKO male mice. (**B**) Representative M-mode echocardiographic images from 10-month-old male WT and ACC dHKO mice. (**C**) Echocardiographic assessment of cardiac function in 10-month-old WT and ACC dHKO male mice. ****P < 0.0001 by unpaired two-tailed Student’s *t* test. (**D**) Total RNA was isolated from hearts of 10-week-old WT and ACC dHKO male mice, and quantitative RT-qPCR was performed to measure expression of hypertrophy-associated genes (*Myh7*, *Nppa, Nppb*). *P < 0.05, ***P < 0.001 by unpaired two-tailed Student’s *t* test. (**E**) Representative trichrome-stained sections from hearts of 11-month-old WT and ACC dHKO male mice are shown at ×10 magnification. Collagen area was quantified from trichrome-positive regions with the split-red-channel method as described in Methods. ****P < 0.0001 by unpaired two-tailed Student’s t test.

Enhanced FA oxidation in the heart has been implicated in impaired cardiac performance (2, 28). Earlier studies in ACC1– or ACC2-deficient mice demonstrated increased FA oxidation specifically in the affected tissues, driven by reduced malonyl-CoA levels. This reduction relieves CPT1 inhibition and enhances mitochondrial FA import and β-oxidation (18, 21, 29–31). Based on these findings, we hypothesized that cardiomyocyte injury in ACC double-knockout (ACC dHKO) hearts results from excessive FA oxidation. To test this, we first assessed CPT1 activity by quantifying long-chain acylcarnitines, products of CPT1-mediated mitochondrial FA transport. Long-chain acylcarnitines were elevated in ACC dHKO hearts, whereas octanoylcarnitine, which enters mitochondria independent of CPT1, was unchanged (Figure 3A).

**Figure 3.**
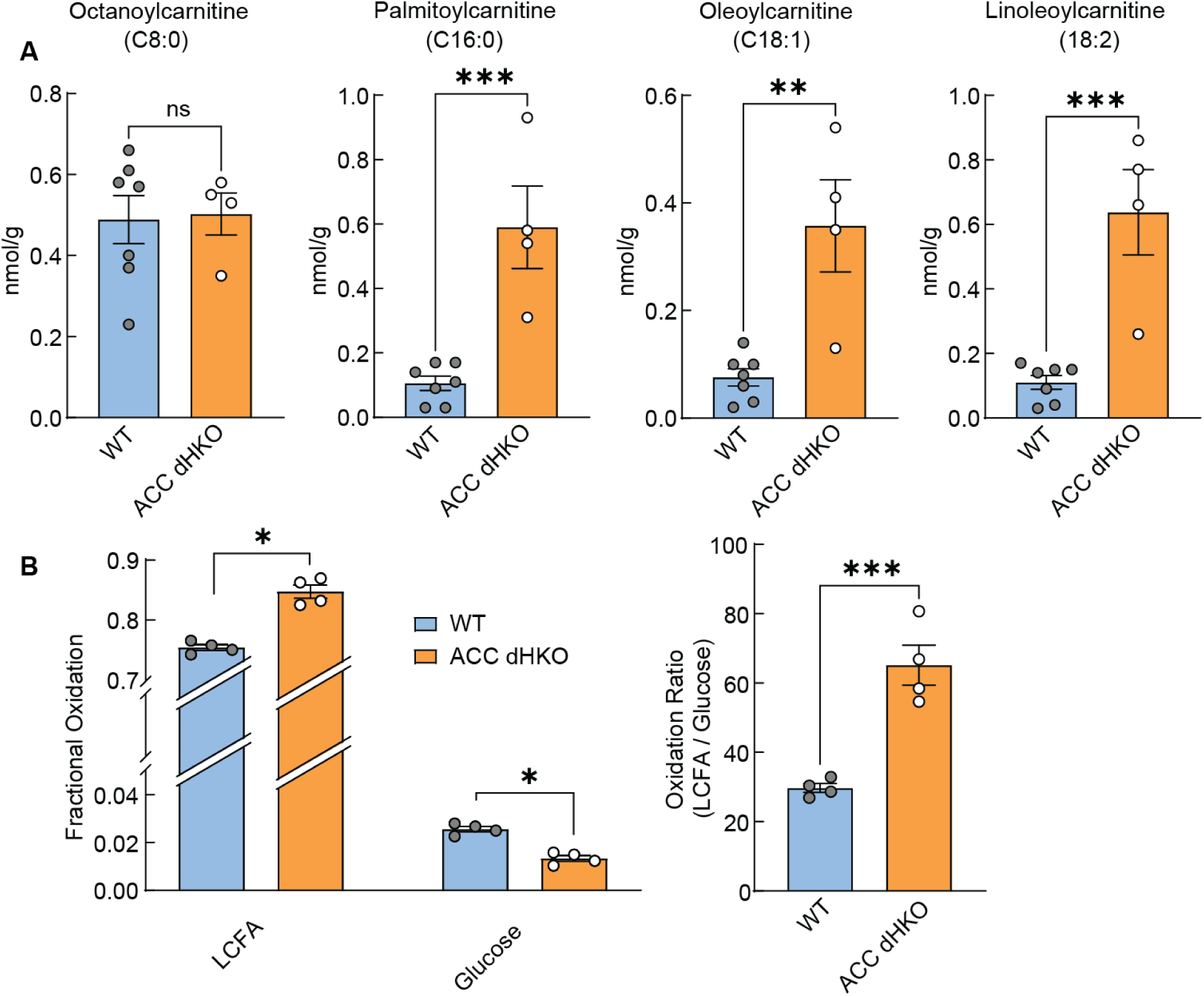
Increased FA oxidation in ACC dHKO hearts. (**A**) Fatty acylcarnitines were measured in hearts isolated from 24-week-old WT and ACC dHKO female mice, as described in Methods. (**B**) Cardiac FA and glucose oxidation rates were measured with Langendorff working heart perfusion in live hearts isolated from 7-week-old WT or ACC dHKO mice (n=4 per group), as described in Methods. Results are shown as the mean ± SEM. *p<0.05, ***p < 0.001, ****p < 0.0001 assessed by Student’s *t* test.

Next, we examined whether FA oxidation was elevated in ACC dHKO hearts due to enhanced CPT1 activity. FA and glucose oxidation rates were measured in isolated Langendorff-perfused WT and ACC dHKO hearts with [U-^13^C] long-chain FA and [1,6-^13^C_2_] glucose tracers. After a 30-min equilibration period to achieve metabolic steady state, fractional substrate oxidation was quantified. ACC dHKO hearts exhibited higher FA oxidation and lower glucose oxidation, leading to an approximately twofold higher FA/glucose oxidation ratio than WT hearts (Figure 3B). These data indicate that ACC deficiency leads to increased CPT1-dependent FA oxidation in the heart.

Given the established link between oxidative stress and mitochondrial dysfunction (32–34), we next assessed mitochondrial integrity in ACC dHKO hearts. Electron microscopy (EM) revealed wider cristae in cardiac mitochondria from ACC dHKO hearts than in WT hearts (Supplemental Figure 1C–D). This cristae dilation is consistent with mitochondrial ultrastructural changes reported in failing hearts and in experimental models of cardiac stress (35, 36). Despite this ultrastructural change, mitochondrial marker proteins and DNA copy number were unchanged between groups, indicating preserved overall mitochondrial content (Supplemental Figure 1E–F).

Next, to assess overall mitochondrial function, mitochondria were isolated and mitochondrial respiratory function was measured with Seahorse (37, 38). ACC dHKO mitochondria exhibited ∼53% lower maximal respiration with complex I substrates (pyruvate/malate) and ∼32% lower respiration with complex II substrates (succinate + rotenone) than WT mitochondria (Figure 4A). In contrast, complex IV–driven respiration (antimycin/TMPD/ascorbate) was unchanged. These data indicate selective impairment of complexes I and II–III in ACC dHKO hearts.

**Figure 4.**
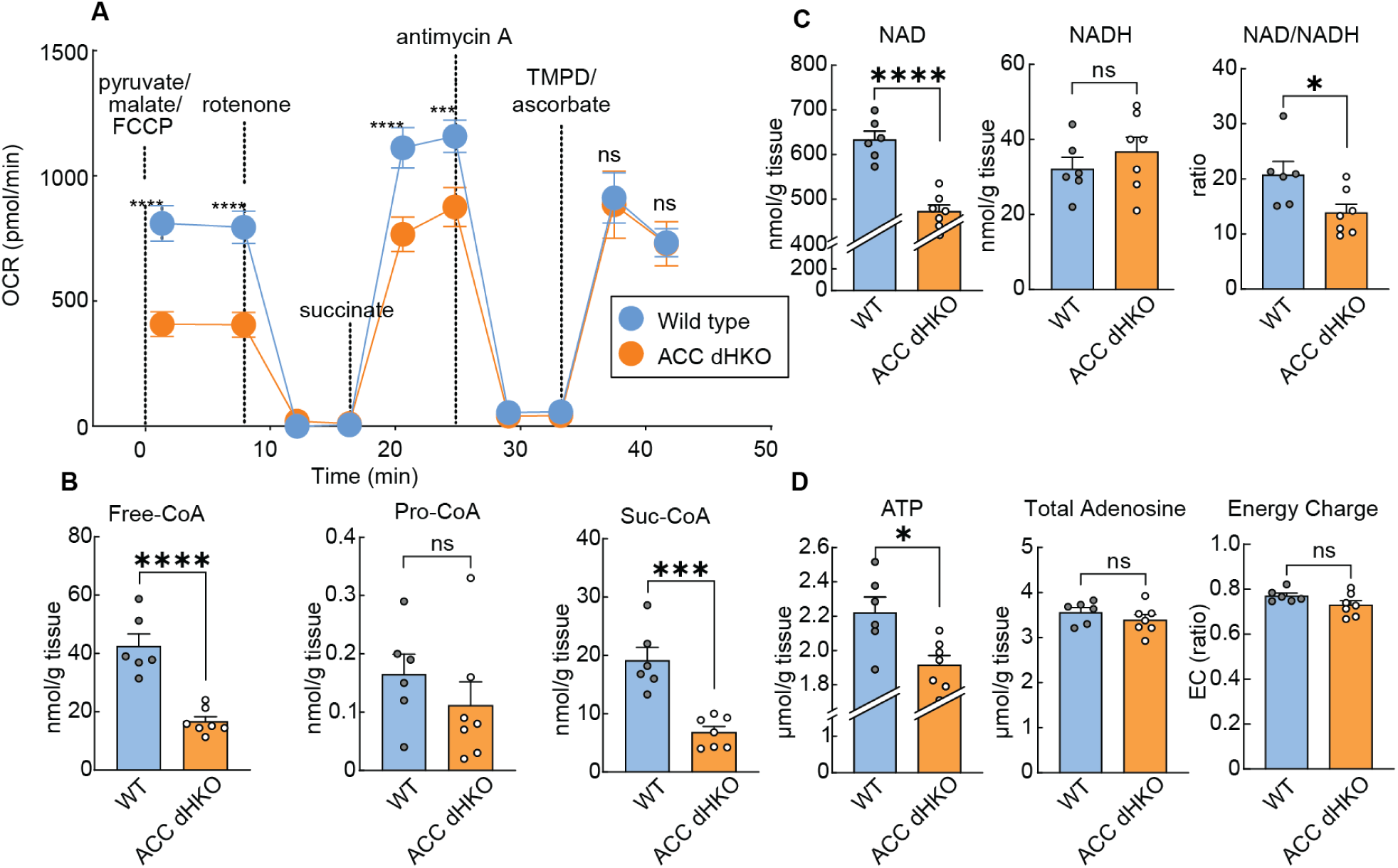
Impaired respiratory electron transfer function in mitochondria isolated from ACC dHKO hearts. (**A**) Mitochondria were isolated from WT and ACC dHKO male mice aged 2–3 months (18 total mice; Seahorse experiments performed in 5 independent runs, 3–4 mice per run with age-matched WT and ACC dHKO in each run). Electron transfer between respiratory complexes was assessed by measuring oxygen consumption rate during sequential additions of pyruvate/malate/FCCP, rotenone, succinate, antimycin A, and TMPD/ascorbate, as described in Methods. (**B**–**D**) Hearts from 20-week-old female WT and ACC dHKO mice were used. (**B**) Free CoA and CoA thioesters (propionyl-CoA and succinyl-CoA) were quantified by LC–MS as described for malonyl-CoA and acetyl-CoA. (**C**) NAD, NADH, and the NAD⁺/NADH ratio were measured by LC–MS in freeze-clamped hearts. (**D**) Total adenine nucleotides levels were quantified by LC–MS. Cellular energy charge was calculated as ([ATP] + 0.5 × [ADP]) / ([ATP] + [ADP] + [AMP]). Data are shown as mean ± SEM. *P < 0.05, ***P < 0.001, ****P < 0.0001 by unpaired two-tailed Student’s *t* test.

To investigate the metabolic basis of this defect, we measured mitochondrial cofactors involved in TCA cycle flux. Free CoA levels were markedly reduced in ACC dHKO hearts, while acetyl-CoA remained unchanged (Figure 4B; see also Figure 1C). Succinyl-CoA was decreased, whereas propionyl-CoA showed a downward trend, consistent with CoA depletion limiting CoA-dependent TCA steps (39, 40).

NAD⁺ and ATP are generated during oxidative metabolism and levels decline in failing hearts (41–43). NAD⁺ and ATP levels in ACC dHKO hearts were reduced by ∼25% and ∼14%, respectively (Figure 4C–D). NADH levels remained unchanged, producing a more reduced NAD⁺/NADH ratio consistent with impaired electron transport. Despite lower ATP, total adenine nucleotides (ATP + ADP + AMP) and cellular energy charge were preserved, indicating maintenance of global adenine nucleotide homeostasis. Together, these findings exhibit that ACC loss in cardiomyocytes leads to reduced essential mitochondrial cofactors, including free CoA and NAD⁺ - constraining TCA cycle flux, redox balance, and oxidative phosphorylation capacity.

We next assessed whether excessive FA oxidation alters cardiac FA composition. ACC dHKO hearts exhibited broad reductions in multiple FAs, including 16:0 (–24%), 18:1n9 (–56%), and 20:4n6 (–34%), with the largest decrease observed in linoleic acid (18:2n6; ∼60%) (Supplemental Figure 2A).

To determine which lipid pools accounted for this loss, cardiac lipids were fractionated into neutral and phospholipid fractions. Neutral lipids (triacylglycerol, diacylglycerol, and cholesteryl esters) and their FA compositions were unchanged (Supplemental Figure 2B–D). In contrast, phospholipids underwent marked remodeling (Figure 5). Although total PE and PG were preserved (Figure 5A–B), total PC and PI were reduced by 36% and 71%, respectively (Figure 5C–D). Importantly, this decrease reflected a selective loss of linoleoyl-containing phospholipid species: most 18:2-containing phospholipids were markedly reduced in ACC dHKO hearts (Figure 5A–D), indicating specific depletion of linoleoyl-phospholipids rather than a global reduction in phospholipid abundance.

**Figure 5.**
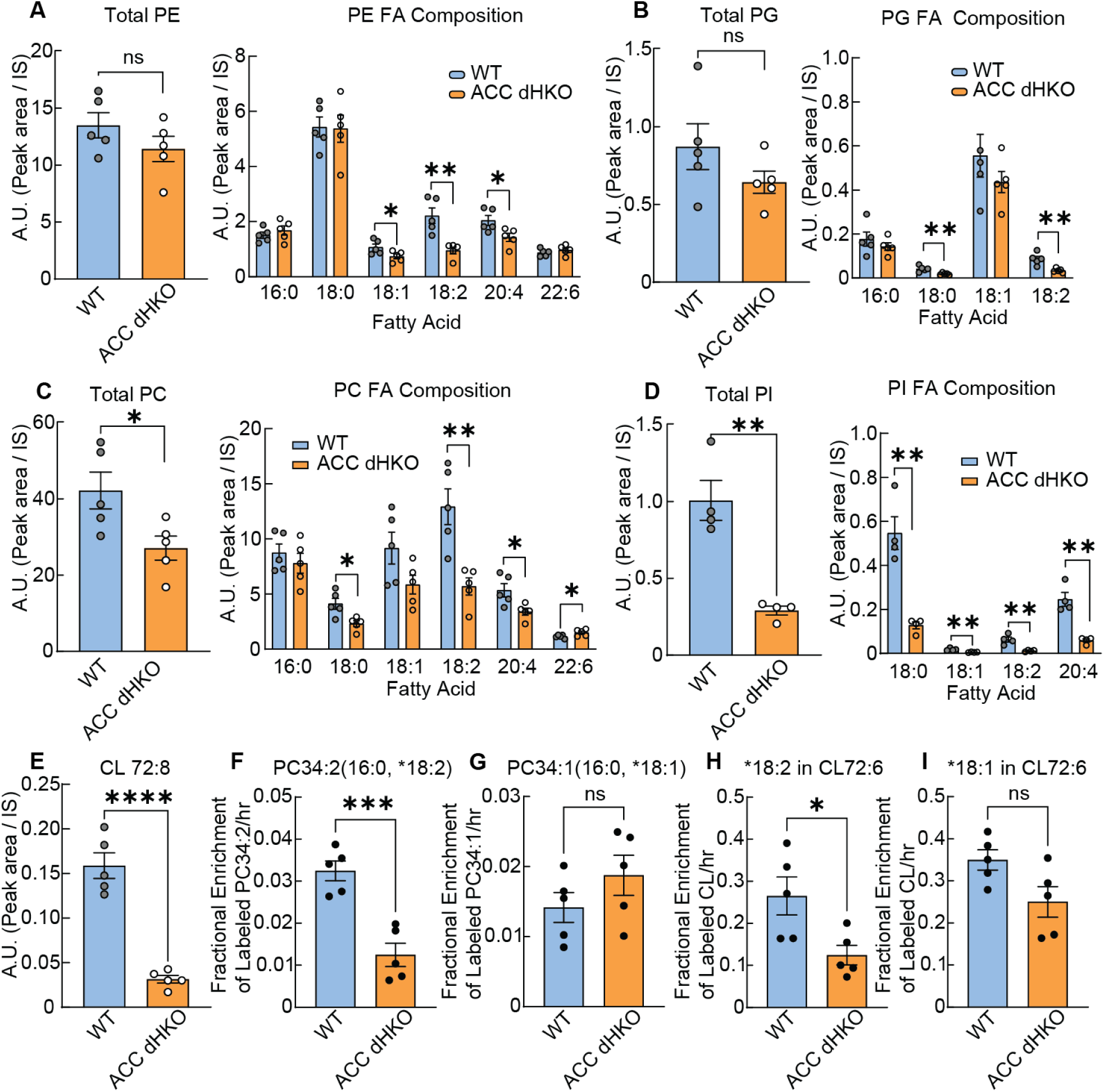
Reduced linoleoyl phospholipids and tetra-linoleoyl cardiolipin in ACC dHKO hearts. (**A**-**E**) Cardiac phospholipids from 8-week-old male WT and ACC dHKO mice were analyzed by LC-MS/MS as described in Methods. Total phospholipids, each PL species, and their specific FA compositions were quantified. PE (**A**), PC (**B**), PI (**C**), PG (**D**), and linoleoyl cardiolipin (72:8, **E**). (**F–I**) Incorporation of labeled FAs into phospholipids and cardiolipins in WT and ACC dHKO hearts. Mice received intraperitoneal injections of LA-d4 (18:2) or OA-d17 (18:1). Enrichment of labeled species was determined one hour post-injection. Mass spectrometry analysis quantified *PC34:2 (16:0,18:2; D4-LA) **(F)**, *PC34:1 (16:0,18:1; D17-OA) **(G)**, *CL72:6 containing one 18:2 chain (D4-LA) **(H)**, and *CL72:6 containing one 18:1 chain (D17-OA) **(I).** Enrichment was calculated as labeled/(labeled + unlabeled) from isotopologue distributions, as described in Methods. Data are presented as mean ± SEM. \**P* < 0.05, \*\**P* < 0.01, \*\*\**P* < 0.001, \*\*\*\**P* < 0.0001 by Student’s *t* test.

Inasmuch as linoleic acid is an essential FA that cannot be synthesized *de novo*, we next determined whether this reduction was due to impaired uptake into cardiomyocytes. To address this, we initially inhibited mitochondrial degradation of infused FAs in ACC dHKO mice by administering the CPT1 inhibitor etomoxir for 40 min and then infused [^3^H]-linoleic acid or [^3^H]-oleic acid for 10 min. The radioactivity was subsequently measured in organic and aqueous phases of heart extracts. No differences in the uptake of 18:2 or 18:1 were measured between WT and ACC dHKO hearts (Supplemental Figure 2E and F).

The marked reduction of linoleic acid–containing phospholipids prompted us to investigate cardiolipin, a mitochondria-specific phospholipid that accounts for ∼20% of total mitochondrial lipids and plays a key role in maintaining respiratory chain function. The predominant mature form of cardiolipin contains four linoleic acid residues (44, 45), and cardiolipin is essential for optimal electron transfer by stabilizing respiratory supercomplexes and supporting efficient oxidative phosphorylation (46–51). Given the close association between cardiolipin composition and mitochondrial respiration, and the abundance of linoleic acid in mature cardiolipin, we measured cardiolipin species in ACC dHKO hearts. Figure 5E shows that tetra-linoleoyl cardiolipin (CL72:8) was ∼80% lower in ACC dHKO hearts than in WT controls, indicating a marked loss of mature cardiolipin. Cardiolipin biosynthesis involves two main steps: the generation of a premature cardiolipin from phosphatidylglycerol (PG) by cardiolipin synthase (CLS), followed by remodeling into the mature form by tafazzin, an acyltransferase that incorporates linoleic acid from 18:2-phosphatidylcholine (PC) or 18:2-phosphatidylethanolamine (PE) (44, 45, 52–54). RT-qPCR analysis revealed that CLS and tafazzin mRNA levels were unchanged in ACC dHKO hearts (Supplemental Figure 2G). In line with this, PG levels were also unaffected (Figure 5B). In contrast, both 18:2-PC and 18:2-PE, which serve as acyl donors for cardiolipin maturation, were markedly decreased by 56% and 55%, respectively, in ACC dHKO hearts (Figure 5A and C).

To determine whether the observed differences in levels of 18:2 containing phospholipids and the mature cardiolipin in ACC dHKO hearts were a result of diminished synthesis, we performed an *in vivo* synthesis assay with isotope-labeled FAs. WT and ACC dHKO mice were administered linoleic acid-d_4_ (LA-d_4_, 18:2) dissolved in saline via intraperitoneal injection. For comparison, oleic acid-d_17_ (OA-d_17_, 18:1) was also injected. One hour later, incorporation of LA-d_4_ into newly synthesized PC34:2 was reduced by 62% in ACC dHKO hearts relative to WT (Figure 5F), whereas incorporation of OA-d_17_ into PC34:1 was unchanged (Figure 5G). Cardiolipin labeling mirrored this pattern: LA-d_4_ incorporation into cardiolipin was reduced by 53% in ACC dHKO hearts, whereas OA-d_17_ incorporation did not differ between groups (Figure 5H–I). These results indicate that the marked loss of tetralinoleoyl cardiolipin (CL72:8) in ACC dHKO hearts is primarily due to insufficient availability of linoleoyl-phospholipid precursors rather than impaired PG supply or reduced expression of cardiolipin biosynthetic enzymes.

Because mitochondrial dysfunction can promote oxidant release, we next assessed oxidative stress. Basal mitochondrial H_2_O_2_ release was slightly elevated in ACC dHKO hearts, though absolute values were low (Supplemental Figure 2H). However, when energized with succinate to drive complex I reverse electron transport (RET), ACC dHKO mitochondria produced markedly less H_2_O_2_ than WT (Supplemental Figure 2I), consistent with reduced electron transport capacity. In line with this, TBARS analysis revealed no increase in lipid peroxidation (Supplemental Figure 2J), and protein carbonylation exhibited only a modest, non-significant trend toward elevation (Supplemental Figure 2K). Thus, ACC dHKO hearts show no evidence of increased oxidative injury, and RET-dependent mitochondrial ROS production is in fact reduced.

PPARα serves as a critical regulator of FA oxidation in the heart, and its activity is reduced proportionally with the decrease of FA oxidation in both animal models and humans with cardiac hypertrophy (55, 56). Downregulation of PPARα is considered the primary mechanism responsible for the shift in substrate utilization from FAs to glucose during the progression of heart failure (2). In hearts of ACC dHKO mice, PPARα signaling was suppressed, similar to findings previously reported in livers of ACC hepatocyte-specific knockout mice (18). In Supplemental Figure 3A, mRNA levels of PPARα-dependent genes were 20-30% lower in hearts from 14-week-old ACC dHKO mice (two left bars for each gene).

To determine whether restoring PPARα activity normalizes the expression of PPARα-regulated genes, the PPAR agonist WY-14643 was administered to 8-week-old ACC dHKO mice for 6 weeks. In Supplemental Figure 3A, WY-14643 restored expression of PPARα-induced genes (*Cpt1a, Cpt1b, Cpt2, HmgCS2*, and *Mcd*) to levels observed in untreated WT hearts. However, despite normalization of PPARα-dependent gene expression, WY-14643 treatment did not improve cardiac function in ACC dHKO mice (Supplemental Figure 3B). Remarkably, early intervention was even detrimental: ACC dHKO mice treated with WY-14643 starting at 4 weeks of age exhibited worse cardiac function after 4 weeks of treatment than those treated beginning at 8 weeks (Supplemental Figure 3C).

To determine whether excessive FAO directly drives the loss of cardiolipin and subsequent heart failure in ACC dHKO hearts, we inhibited FA oxidation with etomoxir. Etomoxir is widely employed in animal studies or cultured cells as an inhibitor of FA oxidation (57, 58). WT and ACC dHKO mice were fed chow supplemented with etomoxir (20 mg/kg/day) for one month, starting at 4 weeks of age. FA composition and cardiolipin levels in the hearts of WT and ACC dHKO mice were then measured. In Figure 6A, etomoxir increased total C18 FAs in the heart, with the largest increase observed in linoleic acid in both WT and ACC dHKO mice.

**Figure 6.**
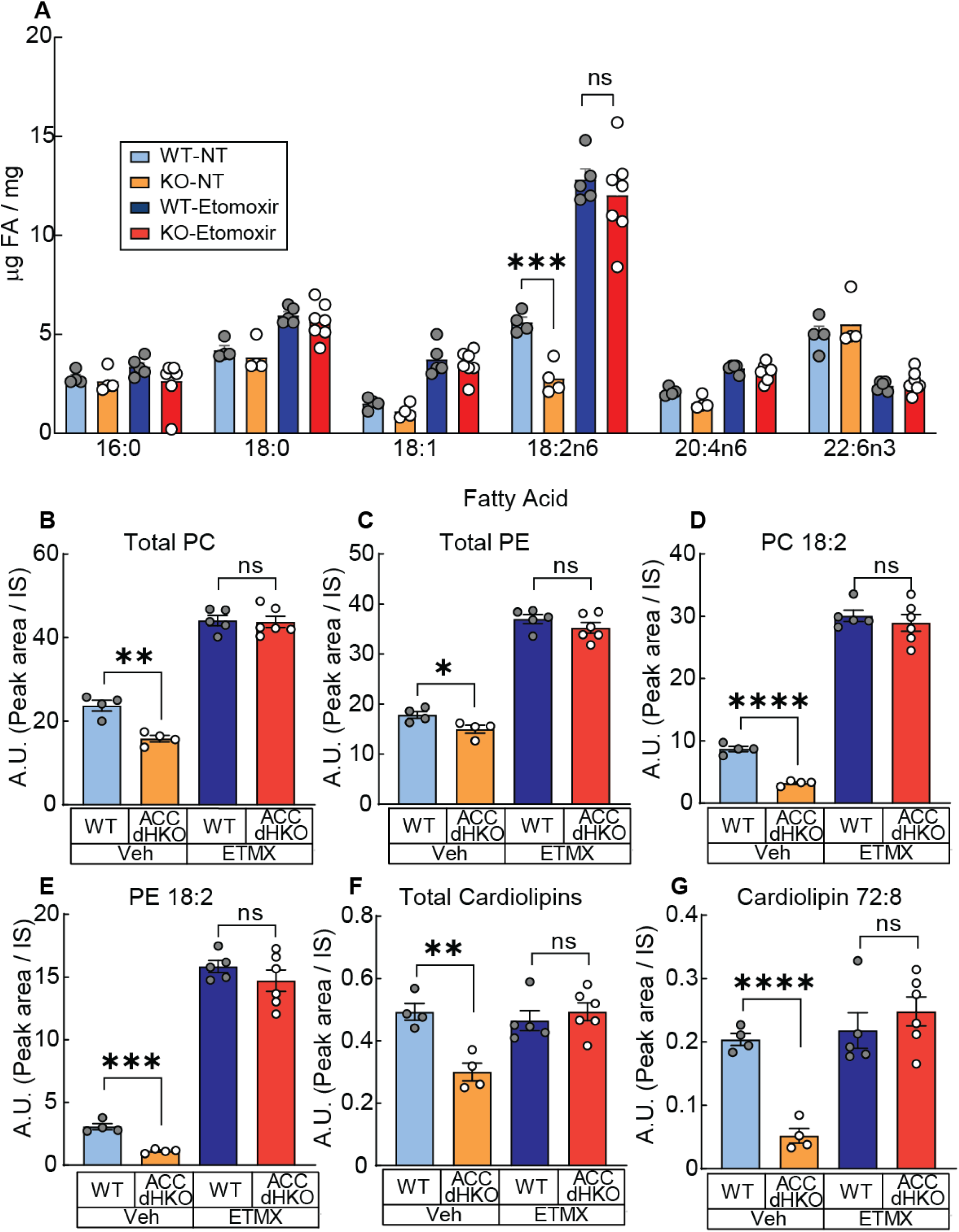
Inhibition of FA oxidation by etomoxir normalizes the amount of linoleic acid in ACC dHKO hearts. (**A**) Hearts from 4-week-old WT and ACC dHKO male mice fed chow or chow supplemented with etomoxir (20 mg kg⁻¹ day⁻¹) for one month were harvested, and FA composition was measured by gas chromatography as described in Methods. (**B-G**) LC–MS/MS was used to quantify total phospholipids (**B**, PC; **C**, PE), linoleoyl-containing phospholipids (**D**, PC 18:2; **E**, PE 18:2), total cardiolipins (**F**), and linoleoyl cardiolipins (**G**, CL 72:8), in hearts from (**A**) as described in Methods. Results are shown as the mean ± SEM. *p<0.05, **p < 0.01, ***p < 0.001, ****p < 0.0001 assessed by Student’s *t* test.

We next analyzed phospholipid species by LC–MS/MS. Etomoxir increased PC and PE by ∼2-fold in both genotypes (Figure 6B–C), and markedly increased linoleoyl-PC and linoleoyl-PE by ∼3-fold and ∼10-fold, respectively (Figure 6D–E). Importantly, this treatment equalized linoleoyl-phospholipid levels between WT and ACC dHKO hearts. Interestingly, total and mature cardiolipin levels were unchanged in WT hearts with etomoxir, but both were restored to WT levels in ACC dHKO hearts (Figure 6F–G). The normalization of linoleic acid concentrations and cardiolipin levels in ACC dHKO hearts following the inhibition of FA oxidation by etomoxir suggests that these changes are primarily driven by excessive FA oxidation.

We next investigated whether etomoxir administration also restores the normal flow of mitochondrial electron transfer through respiratory complexes in the hearts of ACC dHKO mice. WT and ACC dHKO mice, 8 weeks of age, were fed chow supplemented with etomoxir (20 mg/kg/day) for two weeks. The mice were sacrificed for cardiac mitochondria isolation, and the function of electron transport was measured as described in Figure 7A. OCRs were similar across all mitochondrial complexes in mitochondria from WT and ACC dHKO hearts after etomoxir treatment, suggesting that restored cardiolipin levels normalize mitochondrial electron flow in ACC dHKO hearts.

**Figure 7.**
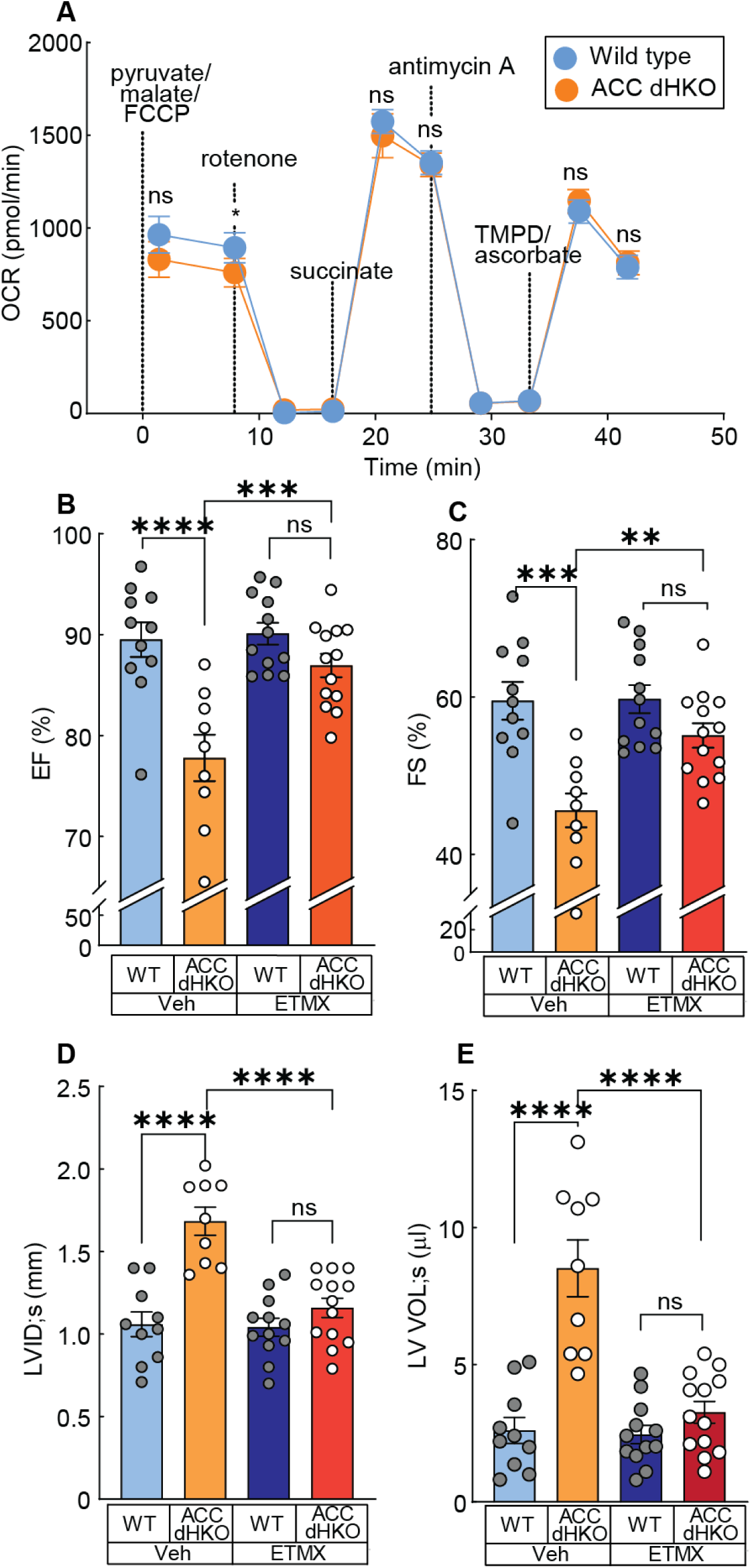
Inhibition of FA oxidation by etomoxir prevents the development of heart failure in ACC dHKO mice. (**A**) Eight-week-old WT and ACC dHKO male mice were fed either chow or chow containing etomoxir (20 mg·kg⁻¹·day⁻¹) for two weeks. Heart mitochondria were isolated, and oxygen consumption rates (OCR) were measured during sequential additions of pyruvate/malate/FCCP, rotenone, succinate, antimycin A, and TMPD/ascorbate, as described in Methods. Data are mean ± SEM; comparisons were made by Student’s *t* test. (**B–E**) Four-week-old WT and ACC dHKO male mice were fed chow or chow supplemented with etomoxir (20 mg·kg⁻¹·day⁻¹) for one month. ECHO was performed to quantify EF; **B**), FS; **C**), LVIDₛ; **D**), and LV Volₛ; **E**). Data shown are mean ± SEM. **p < 0.01, ***p < 0.001 by two-way ANOVA (genotype × treatment) with Sidak’s multiple comparisons test.

Finally, we assessed whether blocking FA oxidation with etomoxir improves the cardiac function of ACC dHKO mice. Mice for these studies are those described for the cardiolipin measurements described above. Chow supplemented with etomoxir (20 mg/kg/day) was fed to 4-week-old WT and ACC dHKO mice for 4 weeks. It is important to note that ACC dHKO mice at 4 weeks of age do not exhibit evidence of heart failure (Supplemental Figure 4A and 4B). As controls, another group of WT and ACC dHKO mice of the same age were fed chow diet. Over the course of one month, chow diet-fed ACC dHKO mice developed statistically significant cardiac dysfunction (Figure 7B-D). Specifically, the EF and FS were reduced by 16% and 28%, respectively, and LVIDs were enlarged by 50% in ACC dHKO hearts. In contrast, hearts from ACC dHKO mice treated with etomoxir displayed normal EF, FS, and LVIDs (Figure 7B-D). These results demonstrate that suppressing FA oxidation with etomoxir effectively prevents the development of heart failure in ACC dHKO mice.

## Discussion

In this study, we demonstrate that loss of ACC1 and ACC2 in cardiomyocytes results in unchecked FAO, depletion of linoleic acid–containing phospholipids, severe cardiolipin deficiency, mitochondrial respiratory failure, and ultimately dilated cardiomyopathy. These findings establish that balanced ACC activity and malonyl-CoA signaling are essential to maintain cardiac lipid composition, mitochondrial function, and contractile performance.

The healthy heart flexibly oxidizes FAs and glucose depending on workload, substrate availability, and hormonal state (28, 59–61). In contrast, failing hearts progressively lose this metabolic flexibility and shift toward greater reliance on glucose oxidation (2, 62). Although the role of FAO in heart failure has been debated, multiple proof-of-concept clinical studies suggest that enhancing glucose oxidation can improve cardiac function. This metabolic shift naturally involves reduced FA oxidation through the Randle cycle (28, 63–67). Consistent with this, direct inhibition of FAO through CPT1 blockade improves cardiac function in myocardial infarction models (58), ischemia–reperfusion model (68), and pacing-induced heart failure models (57), and a pilot trial in patients also reported benefit (16). In agreement with these data, etomoxir treatment fully prevented heart failure in ACC dHKO mice, while oxfenicine, a second compound that inhibits CPT1, produced similar protection and normalized cardiac lipid composition (Figure 7 and Supplemental Figure 4C–G). However, initiating etomoxir treatment after cardiac dysfunction developed (∼20 weeks) did not improve function (Supplemental Figure 5A–F), suggesting that reducing FAO is protective early but does not reverse established disease.

Kolwicz *et al*. (29) reported that cardiomyocyte-specific deletion of ACC2 improved energetic efficiency and protected against pressure-overload-induced dysfunction. Their model reduced malonyl-CoA by ∼50% and modestly increased FAO—conditions that appear adaptive under stress. In contrast, the near-complete loss of malonyl-CoA in our ACC dHKO hearts produced sustained FAO hyperactivation sufficient to cause cardiomyopathy at baseline. Thus, the degree and context of FAO activation determine whether the metabolic shift is beneficial or harmful. Moreover, although *Myh6-*Cre mice can develop late-onset cardiac dysfunction (69, 70), three observations demonstrate that the phenotype here is not due to Cre toxicity: (i) graded severity from ACC1 HKO < ACC2 HKO < ACC dHKO indicates a gene-dosage effect, (ii) etomoxir completely normalized cardiac structure and function, which would not occur with Cre-mediated toxicity, and (iii) abnormalities appeared early in ACC dHKO mice, whereas *Myh6*-Cre effects typically manifest at an old age (70, 71).

Mechanistically, loss of ACC increases FA oxidation and depletes linoleic acid–containing phospholipid precursors, including PC and PE, which are required for the generation of mature tetralinoleoyl cardiolipin (Figure 5A, 5C, 5F–I). In this context, the preferential reduction of multiple FA species in phospholipids—despite preserved levels in neutral lipids such as triacylglycerol, diacylglycerol, and cholesteryl esters—suggests a selective vulnerability of membrane lipids to altered FA flux. Because the heart relies predominantly on lipid oxidation for energy and has limited capacity to store lipids, neutral lipid pools often remain relatively stable even when FA availability or utilization is perturbed (59, 72, 73). By contrast, phospholipids, particularly those comprising mitochondrial membranes, are more sensitive to shifts in FA supply (74). Notably, 18:2 linoleic acid, the dominant acyl chain in cardiolipin, was the most strongly reduced in ACC dHKO hearts, supporting the concept that unrestrained FAO constrains cardiolipin remodeling by limiting linoleate-containing precursor pools.

Given the central role of cardiolipin in organizing the respiratory chain and supporting mitochondrial function, a link between impaired cardiolipin remodeling and mitochondrial dysfunction is well established in Barth syndrome. Cardiolipin is a mitochondrial inner membrane phospholipid that is essential for normal respiratory chain activity (44, 46), and loss of tafazzin—the cardiolipin remodeling enzyme mutated in Barth syndrome—lowers cardiolipin and causes dilated cardiomyopathy in humans as well as in tafazzin-deficient mouse models (75, 76). In these models, cardiolipin deficiency selectively impairs complex I–III function, whereas complex IV activity is relatively preserved (75, 77, 78). Consistent with this paradigm, respiration in mitochondria isolated from ACC dHKO hearts exhibited a pattern similar to that reported in TAZ knockdown or knockout cells (Figure 4A). Together, these observations support cardiolipin deficiency as a major driver of respiratory chain impairment and cardiomyopathy in ACC dHKO hearts, underscoring the requirement for cardiolipin in maintaining complex I–III activity and supercomplex integrity.

Given that disruptions in cardiolipin and respiratory chain organization can secondarily increase ETC-derived oxidative stress, we evaluated whether oxidative damage accompanies cardiolipin loss in ACC dHKO hearts. However, succinate-driven H_2_O_2_ release (reflecting reverse electron transport at complex I) was reduced relative to WT, TBARS levels were unchanged, and the trend toward increased protein carbonylation did not reach statistical significance (Supplemental Figure 2H-K). These findings argue that cardiolipin loss in this model primarily compromises electron transfer efficiency and metabolic balance rather than promoting a generalized increase in oxidative stress.

This framework also helps interpret the stage-dependent shifts in substrate utilization observed in ACC dHKO hearts. The healthy heart adjusts its relative use of FAs and glucose according to workload, substrate availability, and hormonal state, whereas heart failure is often accompanied by progressive loss of metabolic flexibility and a relative shift toward glucose oxidation (2, 28, 62, 79). In ACC dHKO hearts, the FA-to-glucose oxidation ratio was elevated at 7 weeks but markedly reduced at 10 months, when heart failure was established (Figure 3; Supplemental Figure 6A and B). Although reduced FA oxidation at advanced stages of heart failure has been proposed to limit oxygen consumption and oxidative stress and thereby confer functional protection (61, 80–82), an actively regulated metabolic switch is unlikely to account for the late-stage decline in ACC dHKO hearts because malonyl-CoA–mediated control of CPT1 is genetically ablated. Instead, the reduction in FA oxidation appears to be more consistent with a secondary loss of mitochondrial oxidative capacity arising from cardiolipin deficiency and respiratory chain dysfunction.

The selective depletion of linoleate-containing phospholipids in ACC dHKO hearts further raises the question of whether linoleate is lost through oxidative degradation, such as ROS-driven lipid peroxidation, or preferential consumption under sustained FAO. The absence of elevated TBARS and other oxidative damage markers argues against lipid peroxidation as a dominant mechanism. Rather, the data are consistent with a model in which persistently elevated CPT1-dependent FAO promotes preferential β-oxidation of linoleate, thereby depleting linoleoyl-PC/PE pools required for cardiolipin remodeling.

This interpretation is also consistent with the limited efficacy of dietary linoleic acid supplementation. Sunflower oil feeding increased cardiac linoleic acid (C18:2n6) to a similar absolute extent in WT and ACC dHKO hearts (Supplemental Figure 6D), but because baseline linoleate levels were markedly lower in ACC dHKO hearts, this increase was insufficient to restore steady-state linoleate levels and cardiac dysfunction persisted (Supplemental Figure 6E). Thus, under conditions of persistently elevated FAO, increased dietary linoleate can raise cardiac linoleate content but may not achieve the steady-state levels required to support normal phospholipid remodeling, providing a mechanistic explanation for the failure of dietary linoleic acid supplementation to rescue cardiac dysfunction in ACC dHKO hearts.

Along with cardiolipin deficiency, another contributing factor to mitochondrial dysfunction in ACC dHKO hearts may be impaired glucose oxidation. As demonstrated by Szweda *et al*. (83), ACC is required for insulin-stimulated activation of pyruvate dehydrogenase (PDH) via malonyl-CoA–mediated inhibition of FA oxidation. In the absence of ACC, elevated β-oxidation suppresses PDH activity through sustained phosphorylation, leading to decreased glucose oxidation via the TCA cycle. Therefore, the mitochondrial dysfunction observed in ACC dHKO hearts may be compounded by both a deficiency in cardiolipin-dependent electron transport and an impairment in PDH-driven glucose oxidation, further limiting mitochondrial ATP production capacity.

An unexpected observation was the increased docosahexaenoic acid (DHA) levels in ACC dHKO hearts (Supplemental Figure 2A). Sullivan *et al*. (84) reported elevated cardiac DHA in diabetic humans, and DHA supplementation in mice leads to cardiolipin remodeling with increased DHA incorporation, accompanied by reduced respiratory complex activity. Consistent with this, DHA-containing cardiolipins (CL76:12, CL76:11, and CL76:10) were markedly elevated in ACC dHKO hearts (∼5% in WT vs. ∼20% in ACC dHKO; Supplemental Figure 7A and B). Notably, CPT1 inhibition with etomoxir reduced DHA-containing cardiolipin in both WT and ACC dHKO hearts and eliminated the genotype difference (Supplemental Figure 7C), indicating that this remodeling is FAO-dependent. Although we cannot determine from these data whether DHA-containing cardiolipin directly contributes to the decline in cardiac function, its FAO-sensitive regulation and parallel reduction with functional recovery highlight this axis as a potential contributor and a priority for future mechanistic studies.

In addition, analysis of canonical “immature” CL isoforms supports the conclusion that the loss of mature tetralinoleoyl CL (CL72:8) in ACC dHKO hearts reflects impaired maturation/remodeling rather than reduced synthesis of immature CL. These immature species (CL70:X), which serve as tafazzin remodeling substrates (85, 86), were not decreased in ACC dHKO hearts and, in some cases, were modestly increased (Supplemental Figure 7D). Thus, these data support our interpretation that reduced mature CL in ACC dHKO hearts is not driven by diminished immature CL synthesis, but instead reflects a defect in the maturation/remodeling step due to limited availability of linoleate-containing PC and PE.

In summary, cardiomyocyte-specific loss of ACC1 and ACC2 eliminates malonyl-CoA, leading to uncontrolled CPT1-dependent FAO. This sustained FAO depletes tetralinoleoyl cardiolipin—the dominant and functionally essential mitochondrial cardiolipin species—resulting in impaired electron transport, reduced ATP production, and the development of heart failure in ACC dHKO mice. Thus, sustained FAO hyperactivation in this model produces profound energetic and mitochondrial failure, yielding a state of metabolic inefficiency that is conceptually consistent with the broader framework of the failing heart as “an engine running out of fuel (2).” Given the rising burden of cardiomyopathy and limitations of current therapies, our findings highlight the importance of ACC-driven malonyl-CoA signaling and cardiolipin homeostasis in cardiac energy metabolism. Targeting FAO and cardiolipin pathways may offer new therapeutic strategies to prevent or treat heart failure.

## Methods

### Study design and sex as a biological variable

ECHO was performed in male mice because imaging stability and probe positioning were more consistent in males. Biochemical endpoints (including lipidomics, ATP/ADP, and redox assays) were measured in both sexes, and no sex-dependent differences were observed. The study was not powered to detect small sex effects beyond these assays.

### Animals

*Acc1* floxed alleles were generated as we previously described (18). *Acc2^f/f^* mice were purchased from The Jackson Laboratory (stock no. 013042; B6N;129S-*Acacb^tm1.1Lowl^*/J). ACC1 and ACC2 cardiomyocyte-specific double knockout mice were generated by crossing *Acc1^f/f^* and *Acc2^f/f^* mice with *Myh6*-Cre transgenic mice obtained from The Jackson Laboratory (stock no. 011038; B6.FVB-Tg(*Myh6*-cre)2182Mds/J). All animal studies were approved and conducted under the policy of the UT Southwestern Institutional Animal Care and Use Committee. Mice were housed at room temperature (23°C) and maintained on a 12 hr light/dark cycle and provided access to rodent chow diet *ad libitum* (Harlan, Teklad Global 18% Protein Rodent Diet 2018; 18% kcal from fat, 3.1 kcal/g). Etomoxir was purchased from Adooq Bioscience and fed *ad lib* (20 mg/kg/day) after mixing into the powdered chow diet. For echocardiography studies, control flox/flox mice and ACC dHKO mice were fed an etomoxir supplemented diet (20 mg/kg/day) for one month, and for Seahorse studies, mice were fed the etomoxir supplemented diet for two weeks prior to study. A detailed experimental design is provided in the Supplemental Methods.

### RNA extraction and quantitative PCR

RNA was isolated with RNA-Stat60, treated with DNase, and reverse-transcribed with TaqMan reagents. qPCR was performed as described (87). Primer sequences are provided in the Key Resources Table.

### Echocardiography (conscious)

Cardiac function was evaluated by transthoracic ECHO with a VisualSonics Vevo 2100 system equipped with an MS400C probe (88). M-mode images at the papillary muscle level were acquired, and EF, FS, LV dimensions, and wall thickness were calculated per standard guidelines.

### Langendorff heart perfusions

*Ex vivo* substrate utilization was measured in Langendorff-perfused mouse hearts supplied with ^13^C-labeled glucose and FAs. ^13^C-NMR spectra were collected and analyzed by isotopomer modeling to quantify fractional oxidation of each substrate. Experimental conditions, tracers, and analysis parameters are described in the Supplementary Methods.

### FA Composition Measurements

Lipids were extracted from mouse heart with chloroform:methanol (2:1) and dried under nitrogen gas. FA composition was measured with gas chromatography as described previously (89).

### LC-MS/MS lipidomics

Lipids were extracted from ∼50 mg heart tissue with dichloromethane:methanol–based liquid–liquid extraction with internal standards (Splash Lipidomix, Avanti). Samples were analyzed by LC-MS/MS (QTRAP 6500+) and lipid species quantified by MRM and normalized to internal standards. Detailed LC-MS/MS parameters, extraction procedures, and MRM transitions are provided in the Supplementary Methods.

### Mitochondrial respiration

Isolated cardiac mitochondria (5 µg/well) were analyzed with a Seahorse XF24 analyzer. Substrates and inhibitors were injected sequentially (pyruvate/malate/FCCP; rotenone; succinate; antimycin A; TMPD/ascorbate) to assess electron flow across complexes I–IV. Detailed mitochondrial isolation procedures, assay conditions, titration parameters, and sequential injection settings are provided in the Supplementary Methods.

### Short-chain acyl-CoA measurements

Frozen hearts were extracted in 10% trichloroacetic acid spiked with ^13^C-acetyl-CoA and ^13^C-malonyl-CoA, purified by solid-phase extraction, and analyzed by LC-MS/MS in MRM mode. Detailed extraction, SPE conditions, instrument settings, and quantification parameters are provided in the Supplementary Methods.

### Fatty acyl carnitine measurements

Heart tissue was homogenized, spiked with deuterated internal standards, derivatized, separated by reverse-phase HPLC, and analyzed on a triple-quadrupole mass spectrometer (90, 91). Derivatization chemistry, LC gradient, MS transitions, and normalization strategy are detailed in the Supplementary Methods.

### *In vivo* cardiolipin synthesis assays

Mice received intraperitoneal LA-d^4^ and OA-d^17^. Hearts were collected one hour later, lipids were extracted, and LC-MS/MS quantified labeled and unlabeled PC species and cardiolipin. Fractional labeling was calculated as labeled/(labeled + unlabeled). Injection protocol, extraction solvent ratios, MS settings, and isotopologue calculation details are provided in the Supplementary Methods.

### FA uptake assays

Mice were pre-treated with etomoxir (40 mg/kg) for 40 min prior to injection of [^3^H]-linoleic acid or [^3^H]-oleic acid. Hearts were collected 10 min later, extracted by the Folch method, and scintillation counting measured radiolabel incorporation. Tracer preparation, sample handling, phase-separation workflow, and normalization procedures are described in the Supplementary Methods.

### Histology

Heart tissue sections were fixed in 10% (v/v) neutral buffered formalin. Paraffin embedding, sectioning as well as hematoxylin and eosin (H&E) and Trichrome (TC) staining were performed by the UT Southwestern Medical Center’s Molecular Pathology Core. Mean areas of TC-stained collagen in each heart section were determined with standard ImageJ software. TC-stained heart sections demonstrate blue collagen under light microscopy (bright field). The images can be separated into three distinct channels. Collagen in heart sections can be quantified by image reduction to red channel followed by automatic calculation of mean red area (Minimum = 0, Maximum = 255).

## Statistics

Statistical analyses were performed with GraphPad Prism 10. Data are presented as mean ± SEM. Comparisons between two groups were performed with unpaired two-tailed Student’s *t* test. For comparisons among multiple groups, one-way ANOVA followed by Tukey’s post-hoc test was used. *P* < 0.05 was considered statistically significant. The number of biological replicates (*n*) for each experiment is reported in figure legends. No statistical methods were used to predetermine sample size.

## Study approval

All animal experiments were performed in accordance with and approved by the Institutional Animal Care and Use Committee (IACUC) at UT Southwestern Medical Center. No human subjects were included in this study.

## Data availability

Raw data underlying all graphs and reported values presented in the main text and Supplemental figures are provided in the Supporting Data Values file. Lipidomics, LC-MS/MS, isotopic tracing, and Seahorse data are available from the corresponding author upon reasonable request. No high-throughput sequencing or microarray datasets were generated. All materials used in this study are listed in the Supplemental Resource Table.

## Author contributions

C-W.K. and JH conceptualized the study. C-W.K., GV, XF, JM, CD, CL, GS, SD, and MM performed investigation. JH acquired funding. JH, ZW, C.Kh., CM, SC supervised the study. C-W.K., C.Kh., and SD analyzed data. C-W.K. and JH wrote the manuscript. C-W.K., GV, XF, ZW, C.Kh., SC, MM, and JH viewed and edited the manuscript.

## Acknowledgements

We thank Norma Anderson, Judy Sanchez, Tuyet Dang, Tam Tran, and Tessa Edwards for excellent technical assistance, and Chelsea Burroughs and Nancy Heard for assistance with graphics.

