## supplementary materials for "Unrestrained fatty acid oxidation triggers heart failure in mice via cardiolipin loss and mitochondrial dysfunction"

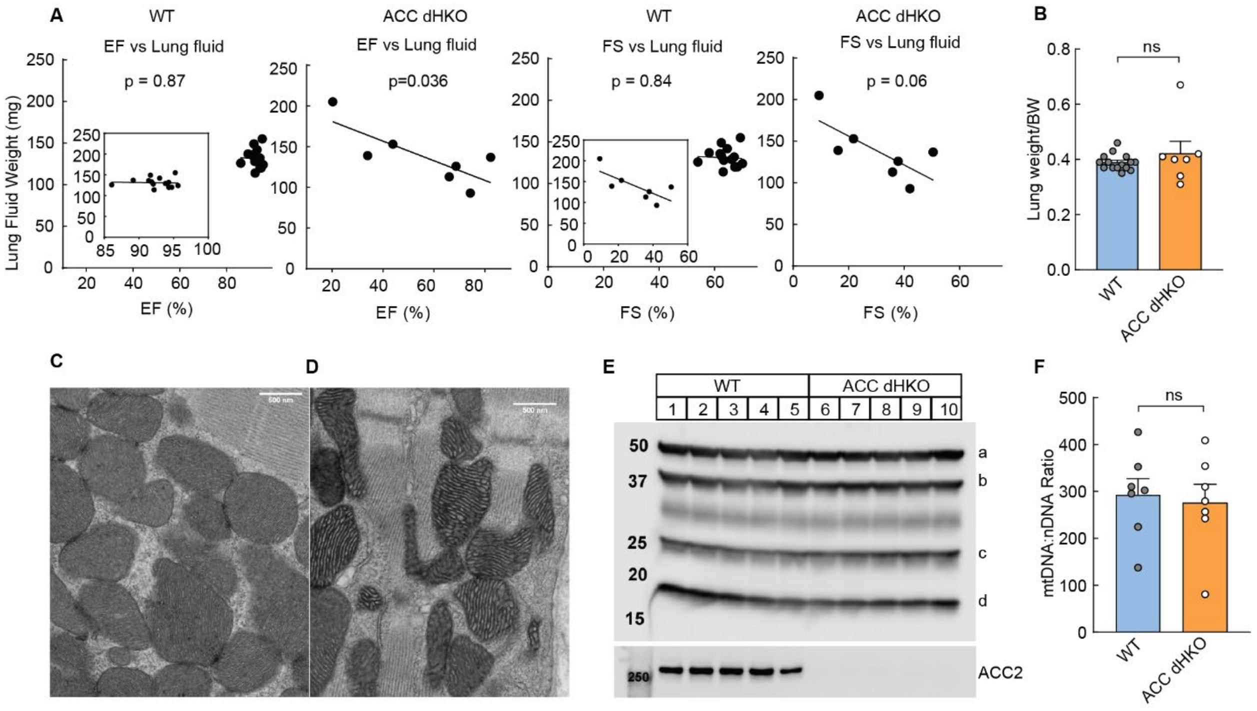

**Supplemental Figure 1.** (**A**) Lung weights from 11-month-old male WT and ACC dHKO mice were measured immediately after euthanasia and drying at 65 °C for 24 h. The correlation between cardiac function measured before euthanasia (EF or FS) and lung weight is shown. Because WT EF/FS values cluster within a narrow range and appear compressed toward the right side of the standardized x-axis, WT data are also displayed as an inset using a magnified x-axis range; the axis labels are the same as in the main panels. (**B**) Lung weight–to–body weight (LW/BW) ratios from 11-month-old male WT and ACC dHKO mice were calculated using lung weights measured immediately after euthanasia and body weights recorded at the time of euthanasia. Data are presented as individual values with mean ± SEM. (**C-D**) Hearts from 6-month-old WT and ACC dHKO mice (2 per group) were analyzed using transmission electron microscopy (TEM) as described in Supplemental Methods (C; WT, D; ACC dHKO). (**E**) Mitochondria isolated from hearts of 20-week-old female WT and ACC dHKO mice and lysates were prepared using RIPA buffer (50 mM Tris pH 7.4, 150 mM NaCl, 1% Triton X-100, 0.5% Sodium deoxycholate, and 0.1% SDS). 30 μg of protein was separated by SDS-PAGE and immunoblot analyses were performed using mitochondrial marker antibodies (a; CV-ATP5A, b; CIII-UQCRC2, c; CII-SDHB, d; CI-NDUFB8). (**F**) Mitochondrial DNA and genomic DNA ratios were calculated as described in Banfi *et al*. (1) . Results are shown as the mean ± SEM. *$p<0.05$, **$p < 0.01$, ***$p < 0.001$, ****$p < 0.0001$ assessed by Student's *t* test.

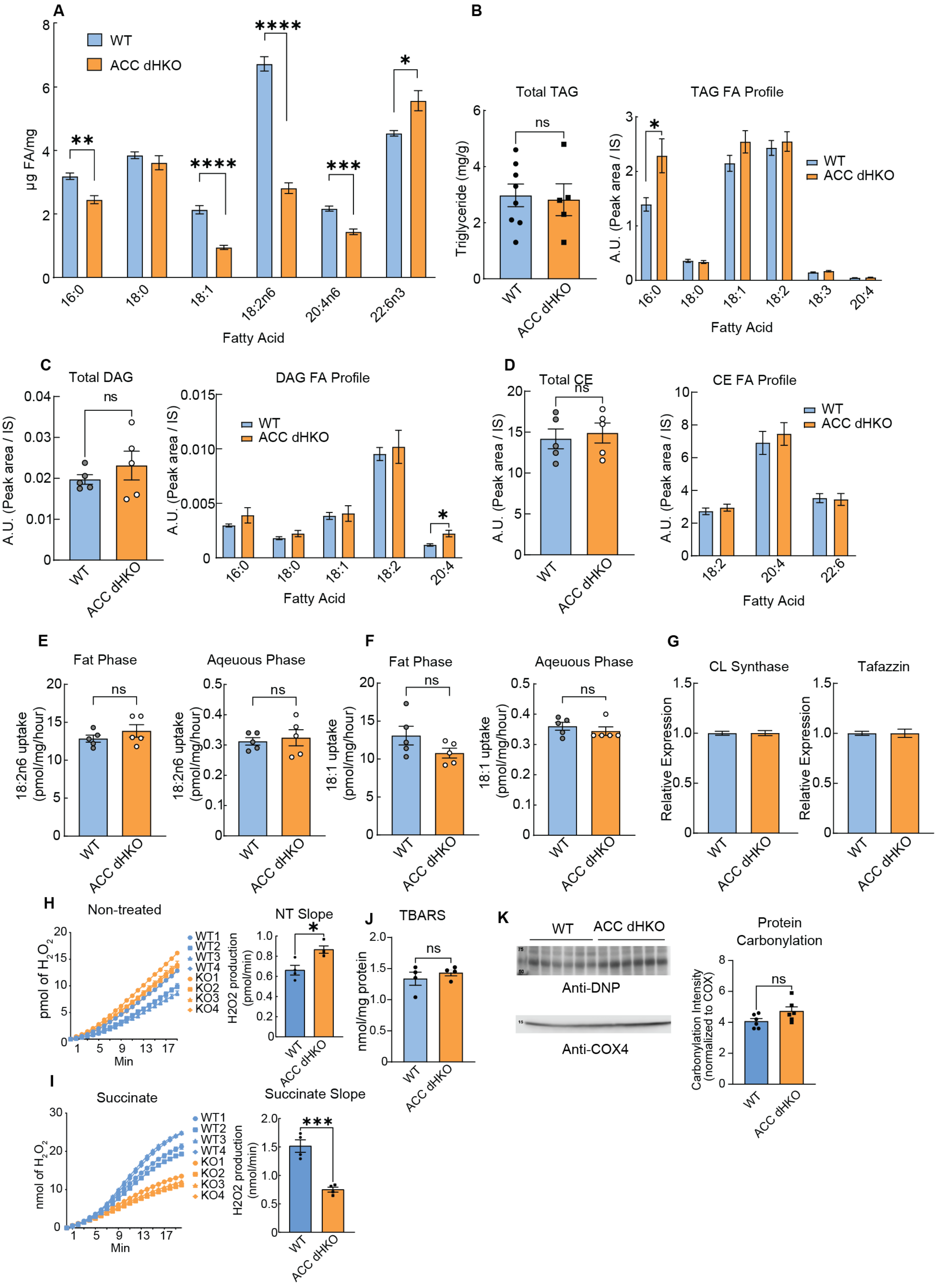
A
WT
ACC dHKO
μg FA/mg
**
****
****
***
*
16:0
18:0
18:1
18:2n6
20:4n6
22:6n3
Fatty Acid
B
Total TAG
ns
Triglyceride (mg/g)
WT
ACC dHKO
TAG FA Profile
*
A.U. (Peak area / IS)
WT
ACC dHKO
16:0
18:0
18:1
18:2
18:3
20:4
Fatty Acid
C
Total DAG
ns
A.U. (Peak area / IS)
WT
ACC dHKO
DAG FA Profile
WT
ACC dHKO
*
16:0
18:0
18:1
18:2
20:4
Fatty Acid
D
Total CE
ns
A.U. (Peak area / IS)
WT
ACC dHKO
CE FA Profile
WT
ACC dHKO
18:2
20:4
22:6
Fatty Acid
E
Fat Phase
Aqeuous Phase
ns
18:2n6 uptake (pmol/mg/hour)
WT
ACC dHKO
F
Fat Phase
Aqeuous Phase
ns
18:1 uptake (pmol/mg/hour)
WT
ACC dHKO
G
CL Synthase
Tafazzin
Relative Expression
WT
ACC dHKO
H
Non-treated
pmol of H2O2
Min
WT1
WT2
WT3
WT4
KO1
KO2
KO3
KO4
NT Slope
*
H2O2 production (pmol/min)
WT
ACC dHKO
I
Succinate
nmol of H2O2
Min
WT1
WT2
WT3
WT4
KO1
KO2
KO3
KO4
Succinate Slope
***
H2O2 production (nmol/min)
WT
ACC dHKO
J
TBARS
ns
nmol/mg protein
WT
ACC dHKO
K
WT
ACC dHKO
Anti-DNP
Anti-COX4
Protein Carbonylation
ns
Carbonylation Intensity (normalized to COX)
WT
ACC dHKO

**Supplemental Figure 2.** (**A**) FA compositions of hearts harvested from 10-week-old WT and ACC dHKO male mice were measured using gas chromatography as described in Methods. (**B**-**D**) Total cardiac triacylglycerol (TAG) was measured biochemically as described in Kim *et al*. (2), diacylglycerol (DAG), and cholesteryl esters (CE) were measured using LC-MS/MS from WT or ACC dHKO hearts as described in Methods. (**E**-**F**) FA uptake (18:1 or 18:2) was assayed in 8-week-old male WT and ACC dHKO hearts as described in Methods. Results are shown as the mean ± SEM. *p<0.05, **p < 0.01, ***p < 0.001, ****p < 0.0001 assessed by Student's *t* test. (**G**) Total RNA isolated from hearts from 10-week-old male WT and ACC dHKO (n=5/group) mice was subjected to qPCR to measure the expression of two major genes in cardiolipin synthesis (cardiolipin synthase and tafazzin). (**H–I**) Heart mitochondria from 20-week-old WT and ACC dHKO male mice were assayed for $H_2O_2$ production using Amplex Red/HRP under basal conditions (**H**) and after succinate addition (**I**). Slopes were calculated from the linear portion of each trace: 6–19 minutes for untreated and 5–14 minutes for succinate-treated samples. (**J**) TBARS levels in whole-heart lysates obtained from 20-week-old female WT and ACC dHKO mice. (**K**) Protein carbonylation in whole-heart lysates obtained from 20-week-old WT and ACC dHKO female mice after DNPH derivatization and anti-DNP immunoblotting; signals normalized to COX4. Data are mean ± SEM. *P* values by Student's *t* test.

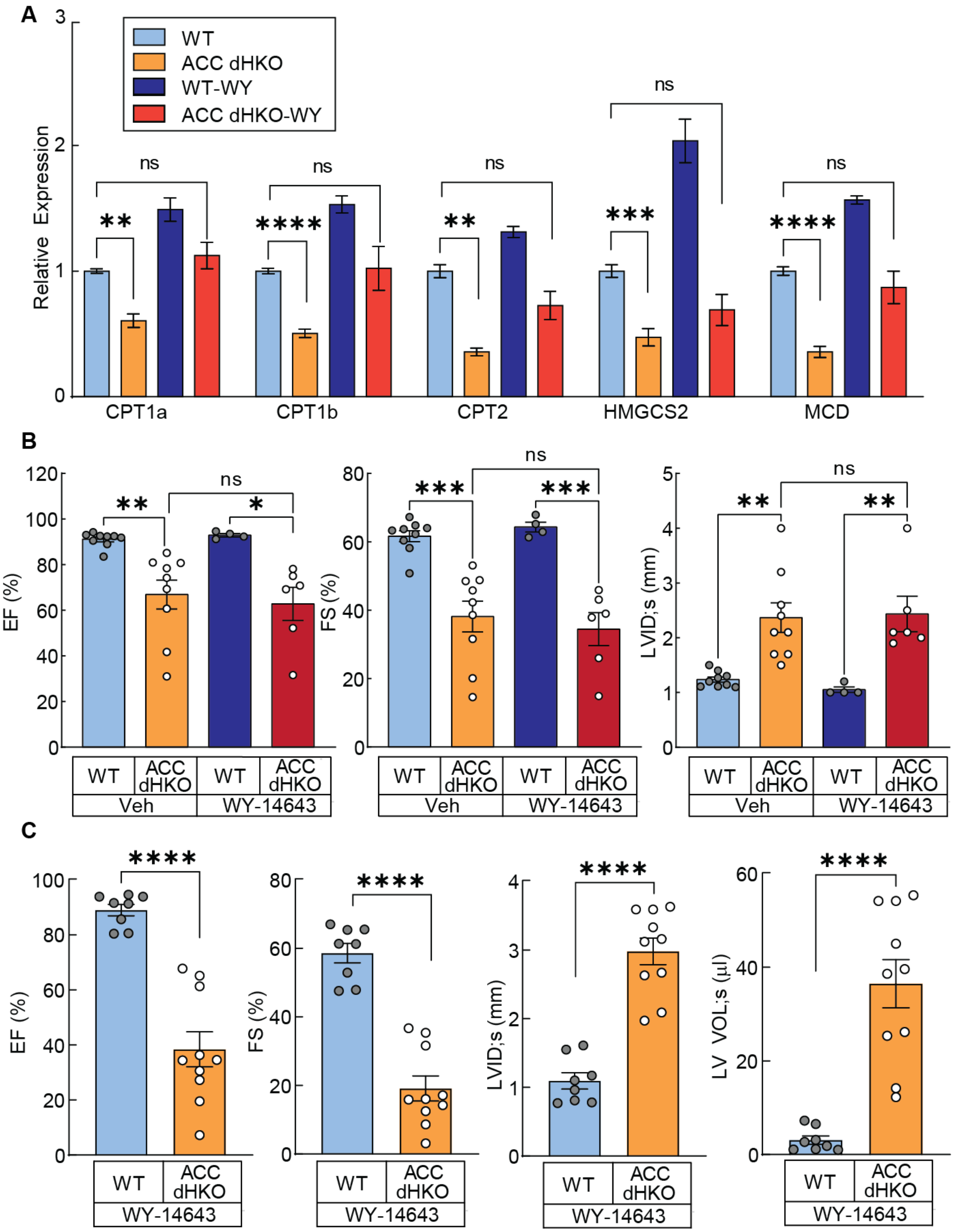

**Supplemental Figure 3.** (**A**) Eight-week-old WT and ACC dHKO mice were fed either chow or chow supplemented with WY14643 (200 ppm) for 6 weeks (n = 4–10 per group). Hearts were harvested, total RNA isolated, and RT-qPCR was performed using primers shown in the figure. (**B**) WT and ACC dHKO mice (12-week-old) were fed either a chow diet or chow supplemented with WY14643 (200 ppm) for 6 weeks (n = 4–10 per group). ECHO analysis was performed to assess the effect of WY14643 on heart function in ACC dHKO mice. Results shown as the mean ± SEM. *p < 0.05, **p < 0.01, ***p < 0.001, ****p < 0.0001 assessed by ANOVA. (**C**) 4-week-old WT or ACC dHKO mice were fed chow or chow supplemented with WY14643 (200 ppm) for 1 month and cardiac function was assessed by ECHO. Results are shown as the mean ± SEM. *p<0.05, **p < 0.01, ***p < 0.001, ****p < 0.0001 assessed by Student's *t* test.

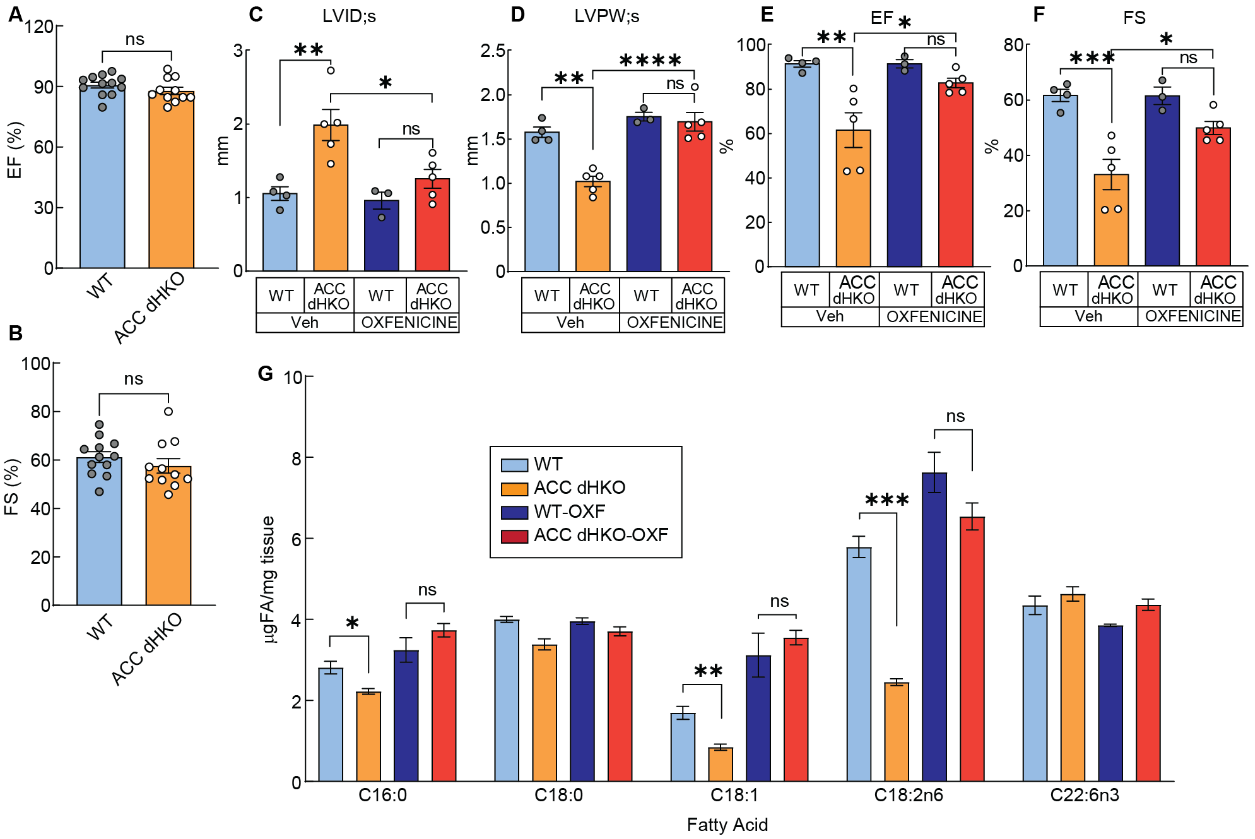

**Supplemental Figure 4.** (**A**-**B**) Cardiac function (EF and FS) of 4-week-old WT or ACC dHKO mice assessed by ECHO. Results are shown as the mean ± SEM assessed by Student's *t* test. (**C**-**F**) Oxfenicine was administered intraperitoneally at 200 mg/kg/day to 8-week-old WT and ACC dHKO mice for two weeks, and cardiac function was assessed by ECHO. (**G**) FA compositions of hearts isolated from WT or ACC dHKO mice treated with Oxfenicine, as described in Figure S4C-F. Results are shown as the mean ± SEM. *p < 0.05, **p < 0.01, ***p < 0.001 assessed by ANOVA.

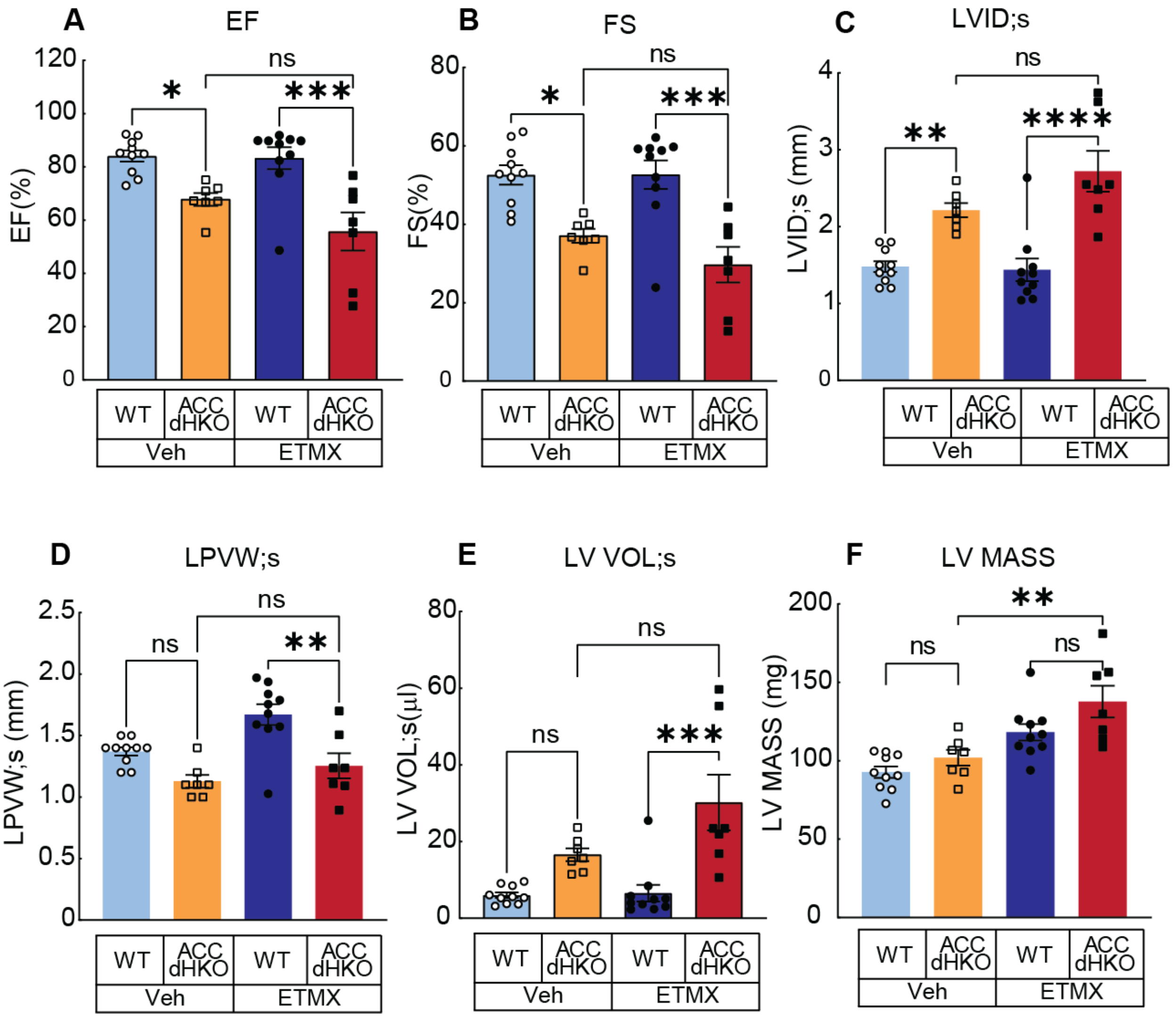

**Supplemental Figure 5.** (**A-F**) Chow supplemented with Etomoxir (20 mg/kg/day) was fed to 20-week-old WT and ACC dHKO mice for one month and cardiac function was assessed by ECHO. Results are shown as the mean ± SEM. *p < 0.05, **p < 0.01, ***p < 0.001 assessed by ANOVA.

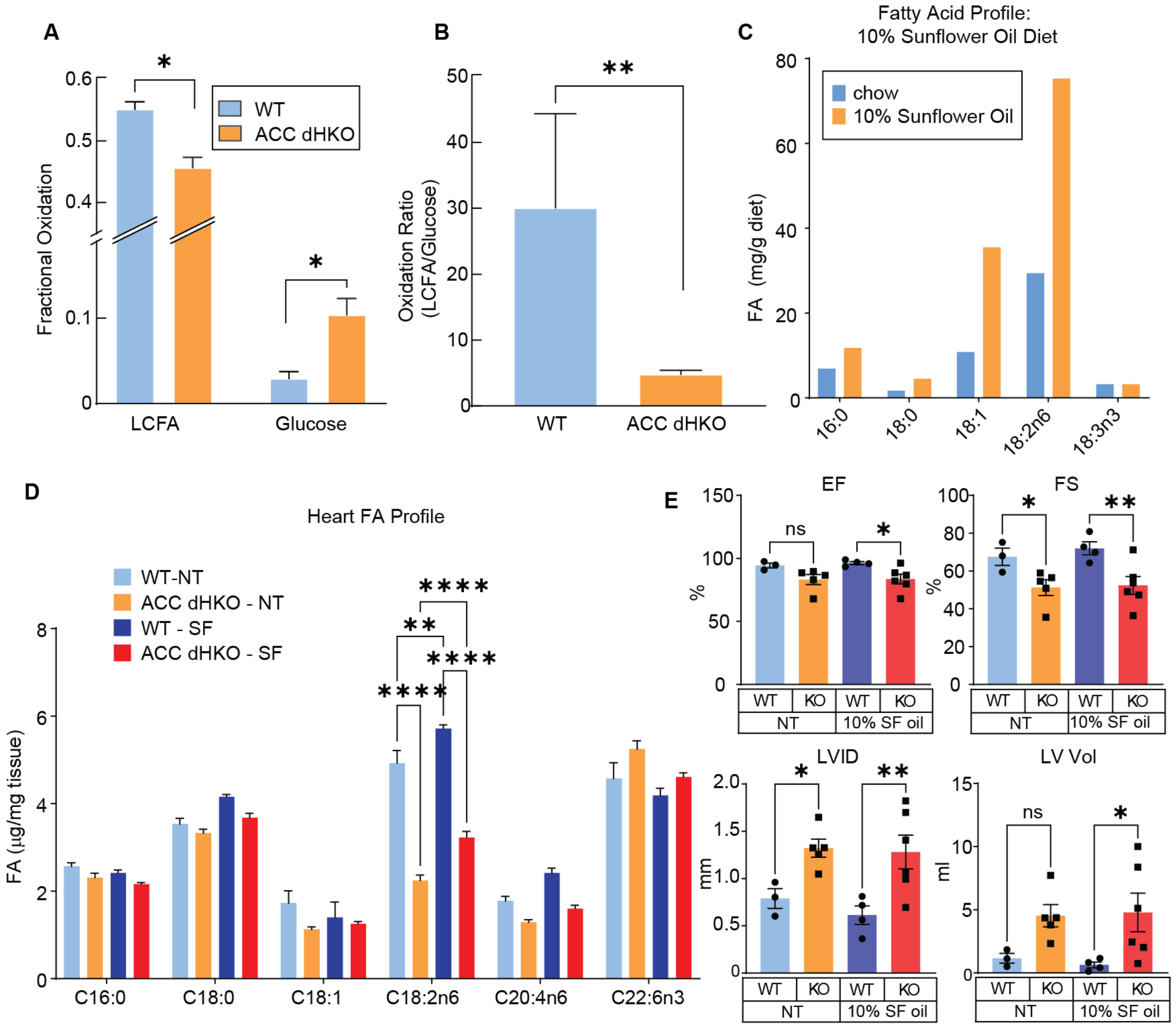

**Supplemental Figure 6.** (**A–B**) FA and glucose oxidation were measured in hearts from 10-month-old WT and ACC dHKO mice (n = 4 per group) using the Langendorff working heart perfusion system, as described in the Methods. (**C–D**) A 10% sunflower oil-supplemented chow diet (based on 2018 regular chow) was fed to 6-week-old female WT and ACC dHKO mice for two months. FA profiles of the diets and mouse hearts were analyzed by GC-FID, as described in the *Methods*. Data are presented as mean ± SEM. *$p < 0.05$, **$p < 0.01$, ***$p < 0.001$ by ANOVA. (**E**) Cardiac function was assessed in mice fed a 10% sunflower oil-supplemented diet for two months, using ECHO, as described in the Methods. Data are presented as mean ± SEM and analyzed by Student’s t test.

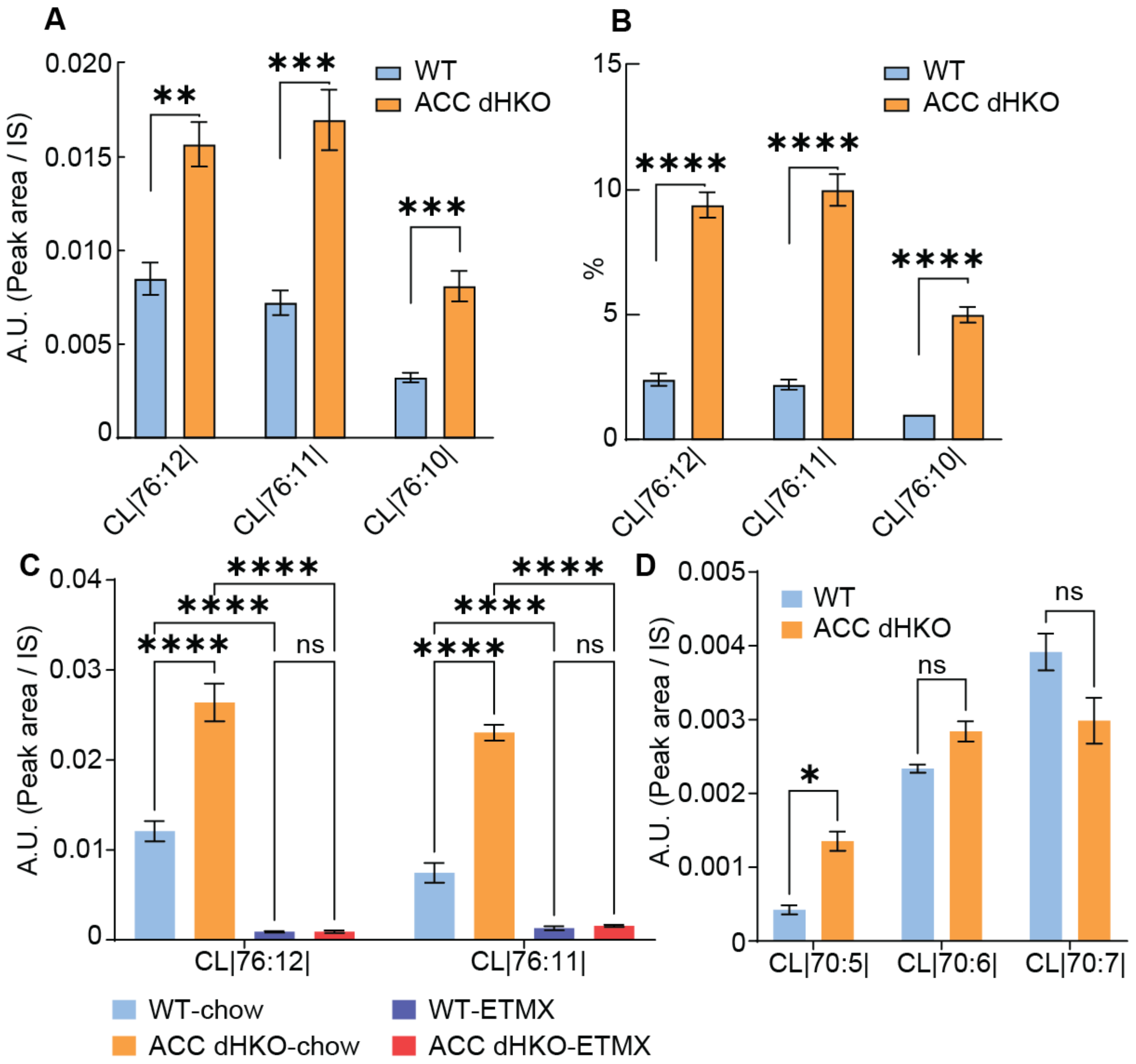

**Supplemental Figure 7.** (**A-B**) Incorporation of docosahexaenoic acid into cardiolipin was measured in hearts from 10-week-old male WT or ACC dHKO (n=5 / group) by LC-MS/MS (A; absolute value, B; Percentage of total cardiolipin) as described in Methods. Results shown as the mean ± SEM. *$p < 0.05$, **$p < 0.01$, ***$p < 0.001$, ****$p < 0.0001$ assessed by ANOVA. (**C**) DHA-containing cardiolipin species were quantified by LC-MS/MS in hearts from WT-chow (n=4), ACC dHKO-chow (n=4), WT+ETMX (n=5), and ACC dHKO+ETMX (n=6) mice. Etomoxir (ETMX; 20 mg $kg^{-1}$ $day^{-1}$) was administered in chow starting at 4 weeks of age for 8 weeks, after which hearts were harvested for lipidomics analysis. Results shown as the mean ± SEM; statistics assessed by ANOVA with multiple comparisons; ns, not significant; ****$p < 0.0001$. (**D**) Representative non–tetralinoleoyl cardiolipin species (CL70:5, CL70:6, and CL70:7) were quantified by LC–MS/MS in hearts from 12-week-old male mice. WT ($n = 4$) and ACC dHKO ($n = 4$) mice were fed chow diet. Peak areas were normalized to internal standards. Statistical significance was assessed by multiple unpaired *t* tests with false discovery rate (FDR) correction. A species was considered significant when the FDR-adjusted q value was < 0.05; nominal p values < 0.05 that did not pass FDR correction are indicated as not significant (ns). Data are shown as mean ± SEM.

### Supplemental Methods and Materials

This file provides expanded protocols corresponding to the brief Methods in the main text; figure/experiment cross-references are noted in each subsection.

#### Transmission Electron Microscopy

This experiment corresponds to Supplemental Figure 1B and C and is not included in the main text. Six-month-old male mice were euthanized and subjected to transcardial perfusion with 4% paraformaldehyde and 1% glutaraldehyde in 0.1 M sodium cacodylate buffer (pH 7.4). After perfusion, hearts were excised and delivered to the Electron Microscopy Core Facility at UT Southwestern Medical Center for further processing. Briefly, the fixed tissues were post-fixed, stained with heavy metals, dehydrated through a graded series of solvents, and embedded in resin. Ultrathin sections were prepared, post-stained, and examined with a JEOL 1400 Plus transmission electron microscope.

#### Measurement of mitochondrial $H_2O_2$ production

This experiment corresponds to Supplemental Figure 2H and I and is not included in the main text. Mitochondrial $H_2O_2$ release was measured using the Amplex Red Hydrogen Peroxide/Peroxidase Assay Kit (Thermo Fisher Scientific) according to the manufacturer’s instructions. Briefly, isolated heart mitochondria were incubated with Amplex Red reagent (10 μM), horseradish peroxidase (1 U/mL), and superoxide dismutase (25 U/mL) in respiration buffer at 37 °C. Fluorescence was monitored at 560 nm excitation and 590 nm emission using a microplate reader, and $H_2O_2$ concentrations were calculated from a standard curve generated with known $H_2O_2$ standards. Substrate conditions included basal, glutamate/malate, and succinate ± rotenone to evaluate forward and reverse electron transport. Non-mitochondrial background signal was subtracted using wells without mitochondria, and catalase addition at the end of each run confirmed assay specificity. Rates were normalized to mitochondrial protein content.

**Protein carbonylation assay**

This experiment corresponds to Supplemental Figure 2K and is not included in the main text. Protein carbonylation was assessed using the Protein Carbonyl Western Blot Detection Kit (ab120419, Abcam) according to the manufacturer's instructions. Briefly, mouse heart tissue was homogenized in ice-cold RIPA buffer containing protease inhibitors, and lysates were clarified by centrifugation. Equal amounts of protein (20 μg) were derivatized with 2,4-dinitrophenylhydrazine (DNPH), neutralized, and resolved by SDS-PAGE, followed by transfer to PVDF membranes. Membranes were blocked and incubated with the anti-DNP antibody supplied in the kit, and immunoreactive bands were visualized by chemiluminescence. Total protein staining served as the loading control. Band intensity was quantified using a LI-COR imaging system and normalized to total protein signal.

**Thiobarbituric acid reactive substances (TBARS) assay**

This experiment corresponds to Supplemental Figure 2J and is not included in the main text. Lipid peroxidation was quantified using the TBARS Assay Kit (Cayman Chemical, Item #10009055) following the manufacturer's instructions with minor modifications. Briefly, ~25 mg heart tissue was homogenized in ice-cold RIPA buffer containing protease inhibitors and centrifuged at 1,600 × g for 10 minutes at 4 °C to obtain clarified lysates. Standards were prepared in the same RIPA buffer. For each reaction, 100 μL lysate was mixed with 100 µL SDS solution and 800 µL TBA color reagent in screw-cap tubes, incubated at 95 °C for 60 minutes, then cooled on ice and centrifuged at 1,600 × g for 10 minutes. A total of 200 µL supernatant per sample was transferred to a 96-well plate and absorbance measured at 530–540 nm using a microplate reader. Malondialdehyde (MDA) concentrations were calculated from an MDA standard curve and normalized to protein content.

**Langendorff heart perfusions**

Utilization of glucose and FA was measured by $^{13}C$-NMR isotopomer analysis in isolated Langendorff perfused mouse hearts procured from WT and ACC dHKO mice. Immediately after cervical dislocation of mice, hearts were excised, arrested in ice-cold saline, cannulated via the aorta, connected to a water-jacketed perfusion column maintained at 37 °C. Hearts were perfused in a retrograde manner for 30 minutes at 100 cm $H_2O$ pressure with a modified Krebs-Henseleit (KH) buffer containing 8 mM [1,6-$^{13}C_2$]glucose, 0.4 mM [U-$^{13}C$]long chain FAs (LCFA), 0.75% bovine serum albumin (BSA). The nonrecirculating buffer was oxygenated with a 95:5 mixture of $O_2$:$CO_2$ using a thin-film oxygenator. Heart rates were monitored throughout the perfusion with a fluid-filled catheter inserted into the left ventricle. Coronary flow samples were collected into a gas-tight syringe at 5 and 25 minutes, followed by measurement of oxygen tension with a blood gas analyzer (Instrumentation Laboratory, Lexington, MA). The difference in oxygen concentrations between the influent and coronary sample, i.e. effluent, was used to calculate myocardial oxygen consumption. At the conclusion of the perfusion, hearts were snap-frozen in liquid nitrogen, pulverized, and extracted with perchloric acid (4%). The extracts were then neutralized and reconstituted in $D_2O$ containing 1 mM ethylenediaminetetraacetic acid (EDTA) and 0.5 mM 2,2-dimethyl-2-silapentane-5-sulfonate (DSS) standard. Proton-decoupled $^{13}$C-NMR spectra of heart extracts were acquired on a 14.1T spectrometer (Bruker Corporation, USA) equipped with a 5-mm cryoprobe. $^{13}$C NMR multiplets from glutamate were deconvoluted using ACD/SpecManager (ACD Labs, Canada). Multiplet ratios for each carbon resonance were used to determine the relative oxidation of [1,6-$^{13}C_2$] glucose, [U-$^{13}$C] FA, and unlabeled endogenous substrates (e.g., triglycerides and glycogen). Multiplet ratios were used to calculate relative fluxes by performing isotopomer analyses using tcaCALC (v.2.07). Data presented as the mean ± SEM (n=4 per group) and analyzed with Welch's *t*-test for statistical significance (* : $P < 0.05$ and ** : $P < 0.01$).

**Sample preparation for LC-MS/MS**

Animal tissues were harvested and immediately snap frozen in liquid nitrogen. The frozen tissues were cut with small disposable razor blades. Approximately 50 mg of heart was weighed and transferred to 2.0 mL prefilled bead beat tubes (2.8 mm ceramic beads). 1 mL of MeOH-DCM (1:2; v/v) was added to the tubes using a serological pipette. Tissues were homogenized in a Bead Ruptor 24 homogenizer from Omni international (Kennesaw, GA, USA) for 50 s (5.5 mps, 3 cycles, 10 s/cycle, 5 s dwell time). The homogenates were transferred to glass tubes and diluted to a final concentration of 10 mg/ml with MeOH-DCM (1:2; v/v). Aliquots corresponding to 0.25 mg of homogenized tissue were transferred to fresh glass tubes for lipid liquid-liquid extraction (LLE).

**LC-MS/MS analysis**

Equal aliquots of mouse heart tissue were transferred to fresh glass tubes for liquid-liquid extraction (LLE). For extraction, 1 mL each of dichloromethane, methanol, and water were added to a glass tube containing the sample. The mixture was vortexed and centrifuged at 2671×g for 5 minutes, resulting in two distinct liquid phases. The organic phase (lower phase) was collected in a fresh glass tube with a Pasteur pipette and spiked with 20 µL of a 1:5 diluted Splash Lipidomix (Avanti Polar Lipids, Alabaster, AL) standard mixture. The samples were dried under $N_2$ and resuspended in 400 µL of hexane.  Lipids were analyzed by LC-MS/MS using a SCIEX QTRAP 6500+ equipped with a Shimadzu LC-30AD (Kyoto, Japan) HPLC system and a 150 × 2.1 mm, 5µm Supelco Ascentis silica column (Bellefonte, PA, USA). The gradient program conditions used are described in table 1. Solvent D (95:5 (v/v) acetonitrile-water with 10 mM ammonium acetate) was infused post-column at 0.03 ml/min. The column oven temperature was 25°C. Data was acquired in positive and negative ionization mode using multiple reaction monitoring (MRM). The LC-MS/MS data was analyzed using MultiQuant software (SCIEX). The identified lipid species were normalized to their corresponding internal standard. Solvents were HPLC or LC/MS grade (Sigma-Aldrich, St. Louis, MO, USA). Splash Lipidomix standards were from Avanti (Alabaster, AL, USA). Lipid extractions were performed in 16×100-mm glass tubes with PTFE-lined caps (Fisher Scientific, Pittsburgh, PA, USA) using glass Pasteur pipettes and solvent-resistant plasticware tips (Mettler-Toledo, Columbus, OH, USA).

Table 1.

| **Time (min)** | **Flow rate (mL/min)** | **% Solvent A*** | **% Solvent B**** | **% Solvent C***** |
|---|---|---|---|---|
| **0** | 0.3 | 97.5 | 2.5 | 0 |

| | | | | |
|---|---|---|---|---|
| **3** | 0.3 | 95 | 5 | 0 |
| **9** | 0.3 | 40 | 60 | 0 |
| **9.3** | 0.3 | 80 | 0 | 20 |
| **20** | 0.3 | 60 | 0 | 40 |
| **26** | 0.3 | 56 | 0 | 44 |
| **26.5** | 0.3 | 40 | 0 | 60 |
| **27.5** | 0.3 | 40 | 0 | 60 |
| **27.6** | 1.2 | 97.5 | 2.5 | 0 |
| **32** | 1.2 | 97.5 | 2.5 | 0 |

*Solvent A – Hexane

**Solvent B – methyl tert-butyl ether (MTBE)

***Solvent C – Isopropanol:Water (90:10, v/v)

**Mitochondrial respiration (electron flow) measurements**

Electron-flow–based mitochondrial respiration was measured using a Seahorse XF24 Extracellular Flux Analyzer (Agilent, Santa Clara, CA, USA) following the manufacturers' instructions. Hearts from WT and ACC dHKO were perfused with 4 mL cold isolation buffer (10 mM MOPS, 1 mM EDTA, 210 mM mannitol, and 70 mM sucrose, pH 7.4). The perfused tissue was disrupted in ice cold isolation buffer using a Potter-Elvehjem homogenizer. Nuclear and unbroken tissues were removed by centrifugation for 10 minutes at 550 x g, at 4°C. The supernatant was filtered through cheesecloth and centrifuged for 15 minutes at 10,000 x g to pellet the mitochondrial fractions. Mitochondria were resuspended in isolation buffer and 5 μg (in 50 ul) was transferred into each well of a XF24 Tissue Plate. Respiration was initially started by incubating isolated mitochondria with 10 mM pyruvate, 2 mM malate, and 4 μM FCCP. Inhibitors or activators of mitochondrial complex were infused to the mitochondria as follows: port A, 50 μL of 20 μM rotenone (2 μM final); port B, 55 μL of 100 mM succinate (10 mM final); port C, 60 μL of 40 μM antimycin A (4 μM final); port D, 65 μL of 100 mM ascorbate plus 1 mM TMPD (10 mM and 100 μM final, respectively). $O_2$ consumption (OCR) was measured at intervals of 4 - 5 minutes (3).

**Short-chain acyl-CoA quantification**

Approximately 50 mg of freeze-clamp frozen heart samples were homogenized in 500 μl ice-cold 10% trichloroacetic acid and immediately spiked with labeled [$^{13}C_2$]acetyl-CoA and [$^{13}C_3$] malonyl-CoA for an internal standard. Samples were centrifuged at 4°C for 10 minutes and 150 μL supernatant was loaded onto an Oasis HLB 1cc-30 mg solid-phase extraction column. The column was washed with water and methanol. The eluent was lyophilized and dissolved in 50 μL of mobile phase of which 1 μl was subjected to LC/MS/MS analyses.

Analysis was performed on an API 3200 triple quadrupole LC/MS/MS mass spectrometer (Applied Biosystems/Sciex Instruments) in positive electrospray ionization mode. The mass spectrometer was equipped with a Shimadzu LC-20AD liquid chromatograph (LC) and a SIL-20ACHT auto sampler. Chromatography was performed on a reverse-phase C18 column (Waters xBridge, 150 x 2.1 mm, 3 μm) with a mobile phase consisting of water/methanol (95:5, v/v) and 4 mM dibutylamine acetate (eluent A), and water/acetonitrile (25:75, v/v) (eluent B). MRM was carried out for the detection of the short-chain acyl-CoAs in standard solutions and biological samples. Acetyl-CoA and malonyl-CoA were quantified by comparing the individual ion peak area with that of the internal standard. The analytical data was processed by Analyst Software (version 1.6.2).

**Fatty acyl carnitine measurements**

Fatty acyl carnitines were measured as described in Millington *et al.* (4). Approximately 20 mg of snap-frozen heart tissue was homogenized in water. The samples were then centrifuged at 16000 x g, 4 °C, for 10 minutes. The supernatant from the centrifugation was spiked with deuterium-labeled acylcarnitine standards (Cambridge Isotope Laboratories, Andover, MA) and proteins were removed by precipitation with acetonitrile. The derivatization and measurements were performed as described previously (5). Chromatographic separation was achieved on a C18 column (Xbridge, Waters, Milford, MA; 150×2.1 mm, 3.0 μm). The analysis was performed in MRM mode on API 3200 triple quadrupole LC/MS/MS mass spectrometer with electrospray ionization. Quantification of acylcarnitines was achieved by comparison of the individual ion peak area with that of the internal standard.

***In vivo* cardiolipin synthesis assay**

1 mg/kg of d17-labeled oleic acid (18:1) and d4-labeled linoleic acid (18:2) were administered to WT or ACC dHKO mice via intraperitoneal injection. The mice were euthanized exactly one hour later, and the hearts were collected. Samples were extracted by a modified Bligh-Dyer extraction. Briefly, 50 mg of heart tissue was homogenized with a bead beater in 1 mL methanol (MeOH) and a recovery standard of d9-PC34:2 (Avanti Polar Lipids) was added. After homogenization, the samples were diluted to a final concentration of 1:1:2 ratio of water, MeOH, and dichloromethane to a total volume of 8

mL. After centrifugation, the bottom phase was removed and dried under nitrogen. The sample was resolubilized in 1 mL 90% MeOH, 10% water. The processed samples were analyzed on an LC-MS/MS system (Shimadzu LC20 paired with Thermo Fusion Lumos) targeting CL72:6(18:1_18:1_18:2_18:2), PC34:2(16:0_18:2) and PC34:1(16:0_18:1). The M+0 (m/z 281.2 and m/z 279.2), M+4(m/z 285.2 and m/z 283.2), and M+17 (m/z 299.2 and m/z 297.2) peaks were monitored. The samples were separated on poroshell C18 column (Agilent) with a linear gradient transitioning from 90% MeOH/10% water to 100% MeOH over 28 minutes. The lipids were fragmented in negative mode for both the labeled and unlabeled FAs (M+0 for unlabeled, M+4 for linoleic acid, and M+17 for oleic acid). The reported fraction labeled is the ratio of the peak area of the labeled FA to the sum of both the labeled and unlabeled peak of each FA in each lipid.

**Measurement of FA Uptake in Heart**

To assess FA uptake into the heart, 10 mice per group were injected intraperitoneally with etomoxir (40 mg/kg, 2.5 mg/mL in saline) to inhibit FA oxidation. After 40 minutes, each mouse received 5 µCi of tritium-labeled linoleic acid ($^3$H-LA) or oleic acid ($^3$H-OA) via retro-orbital injection in 200 µL of 0.05 mM albumin-conjugated FA containing 0.1 mM BSA. Ten minutes later, mice were euthanized with isoflurane, and hearts were collected, rinsed in PBS, and weighed. Tissues were homogenized in 1 mL of Folch reagent (chloroform:methanol, 2:1) with ceramic beads, followed by centrifugation at 10,000 rpm for 5 minutes at 4°C. The supernatant was re-extracted with 1 mL of fresh Folch reagent, and the pooled extracts were mixed with 1 mL of 1 M $CaCl_2$, vortexed, and centrifuged at 3,000 rpm for 10 minutes to separate aqueous and organic phases. One milliliter of each phase was transferred to scintillation vials, dried, and mixed with scintillation cocktail for radioactivity measurement. Uptake was normalized to blood radioactivity measured from 50 µL of whole blood.

**Resource Table**

| REAGENT or RESOURCE | SOURCE | IDENTIFIER |
|---|---|---|
| **Antibodies** | | |
| Rabbit anti-ACC1 | Kim et al., (2) | |
| Total OXPHOS Rodent WB antibody Cocktail | Abcam | Cat#ab110413 |
| **Chemicals** | | |
| Wy 14643 – 250 mg | Cayman Chemical | Cat#70730 |
| Seahorse XF24 Islet Capture Microplates | 101122-100 | Agilent |
| 4-Hydroxy-L-phenylglycine (Oxfenicine) | Sigma | Cat#56160-50G |
| Etomoxir | AdooQ Bioscience | Cat#A11415 |
| 1-(2,3,4-Trimethoxybenzyl)piperazine dihydrochloride (TMZ) | Sigma | Cat#653322-10G |
| Oleoyl-L-carnitine | Cayman | Cat#26557-5mg |
| Linoleoyl-L-carnitine (chloride) | Cayman | Cat#26560-1mg |
| L-Glutamic acid | Sigma | Cat#G8415-100G |
| L-(−)-Malic acid | Sigma | Cat#M6413-25G |

| | | |
|---|---|---|
| Sodium succinate dibasic hexahydrate | Sigma | Cat#S2378-100G |
| Rotenone | Sigma | Cat#R8875-1G |
| Oligomycin | Sigma | Cat#O4876-5MG |
| FCCP | Sigma | Cat#C2920-10MG |
| Antimycin A | Sigma | Cat#A8674-25MG |
| N,N,N′,N′-Tetramethyl-p-phenylenediamine | Sigma | Cat#T7394-5g |
| L-Ascorbic acid | Sigma | Cat#A5960-25G |
| Oleoyl-L-carnitine-d3 (chloride) | Cayman | Cat#26578-1mg |
| Pyruvic acid | Sigma | Cat#107360-100G |
| LINOLEIC ACID [9,10,12,13-3H] | ARC | ART 0332-250 µCi |
| Palmitic Acid, [9,10-3H(N)]-, 5mCi (185MBq) | PerkinElmer | NET043005MC |
| Oleic acid, [9-10-3H(N)] | ARC | ARC 09182 – 5 mCi |
| **Critical commercial assays** | | |
| RNA STAT-60 kit | TEL TEST | Cat#NC9489785 |
| DNA-free, DNA removal kit | Invitrogen | Cat#1906 |

| | | |
|---|---|---|
| Taqman reverse transcription reagents | Applied Biosystems | Cat#N8080234 |
| 2× SYBR Green PCR master Mix | Applied Biosystems | Cat#4309155 |
| SuperSignal West Pico Chemiluminescent Substrate | Thermo Fisher Scientific | Cat#34580 |
| Pierce BCA Protein Assay Kit | Thermo Fisher Scientific | Cat#23225 |
| TBARS Assay Kit | Cayman | Cat#10009055 |
| Amplex™ Red Hydrogen peroxide/Peroxidase Assay Kit | Thermo Fisher Scientific | Cat#A22188 |
| Protein Carbonylation Kit | Abcam | Cat#ab120419 |
| **Primers for quantitative real-time PCR** | | |
| *ACACA (Acc-1)* | | |
| 5' primer: 5'- CACGGGCAGTCTACCACAGA-3'<br>3' primer: 5'- AGTGGAAAACTCGATGGAGCTT-3' | | |
| *ACACB (Acc-2)* | | |
| 5' primer: 5'-GCTGCCGACGGGATCAG-3'<br>3' primer: 5'-TCCAGACACATTGAGCATGTCAT-3' | | |
| *Tafazzin* | | |
| 5' primer: 5'- CCCTCCATGTGAAGTGGCCATTCC -3'<br>3' primer: 5'- TGGTGGTTGGAGACGGTGATAAGG -3' | | |

| |
|---|
| *Cardiolipin synthase* |
| 5’ primer: 5’- GGTGTTGCACAGCATTCA -3’<br><br>3’ primer: 5’- GCTGGATCTGGGTGCTTCT -3’ |
| *Cpt-1a* |
| 5’ primer: 5’- CACCAACGGGCTCATCTTCTA -3’<br><br>3’ primer: 5’- CAAAATGACCTAGCCTTCTATCGAA -3’ |
| *Cpt-1b* |
| 5’ primer: 5’- GCACTTCTCAGCATGGTCATCT -3’<br><br>3’ primer: 5’- GGGTTTGTCGGAAGAAGAAAATG -3’ |
| *Cpt2* |
| 5’ primer: 5’- AGCCTACCTGGTCAATGCATATC -3’<br><br>3’ primer: 5’- GGGTTTGGGTATACGAGTTGAATT -3’ |
| *Hmgcs2* |
| 5’ primer: 5’- GAGTTAAGGCACCTGCTACTAACCTT -3’<br><br>3’ primer: 5’- GACACTTTCAGGAATGGGTTATCTC -3’ |
| *Mlycd* |
| 5’ primer: 5’- GGAGACAGGCCCCAACAGT -3’<br><br>3’ primer: 5’- TGAGGATCTGCTCGGAAGCT -3’ |
| *Pdk-4* |
| 5’ primer: 5’- AAGCAAAACACAAACACGAGTACAA -3’<br><br>3’ primer: 5’- CCCGGGTCATCCAACCA -3’ |
| *Hmgsyn* |
| 5’ primer: 5’- GCCGTGAACTGGGTCGAA -3’ |

| | | |
|---|---|---|
| 3’ primer: 5’- GCATATATAGCAATGTCTCCTGCAA -3’ | | |
| **Software and algorithms** | | |
| Image Studio™ Version 5.0 | Li-COR | N/A |
| Prism 9 | GraphPad | https://www.graphpad.com/ |
